# The lipid A acylation pattern of *Coxiella burnetii* prevents detection and clearance by the non-canonical inflammasome in primary murine macrophages

**DOI:** 10.64898/2026.05.07.723481

**Authors:** Manuela Szperlinski, Faiza Asghar, František Csicsay, Elias Schermuly, Roland Lang, Ludovit Skultéty, Christian Berens, Katja Mertens-Scholz, Anja Lührmann

## Abstract

*C. burnetii* is a Gram-negative, obligate intracellular bacterium and the causative agent of Q fever. The disease is either asymptomatic or manifests as a mild flu-like illness, but pneumonia or hepatitis might also occur. In most cases, the infection is self-limiting and the pathogen is cleared. In a small percentage of patients, the host immune system fails to eliminate the pathogen, potentially allowing the development of chronic Q fever months or even years after primary infection. The elimination of the bacteria, and thereby prevention of disease onset, would require an inflammatory response. Inflammasomes are multimeric protein complexes that induce a pro-inflammatory response to combat pathogens. Here we show that *C. burnetii* fails to induce a strong activation of the non-canonical inflammasome, independently of its type IVB secretion system. However, the pathogen is unable to prevent external activation of the non-canonical inflammasome, which subsequently results in a reduction of the bacterial burden. Importantly, the acylation pattern of lipid A was identified to be involved in avoiding the activation of the non-canonical inflammasome. *C. burnetii* harbors a tetra-acylated lipid A. Modification of the *C. burnetii* lipid A to penta-/hexa-acylation resulted in increased secretion of IL1β and reduced bacterial load. Together, these results suggest that the acylation pattern of lipid A constitutes an important immune evasion strategy of *C. burnetii* by failing to activate the non-canonical inflammasome. In addition, evidence was provided that oxygen limitation arrests activation of the NLRP3 inflammasome in murine BMDM, which might prevent efficient elimination of bacteria under hypoxic conditions, such as in granulomas or in inflamed tissue.

**AUTHOR SUMMARY:** Several pathogens have evolved mechanisms to persist in the human host, which allows reoccurring or late onset of infection. The human innate immune system has therefore established several pathways, including the inflammasome, to prevent bacterial survival. Here we show that the obligate intracellular pathogen *Coxiella burnetii*, the causative agent of Q fever, prevents detection by the non-canonical NLRP3 inflammasome. This is mediated by the acylation pattern of its lipid A. Altering this acylation pattern allows activation of the inflammasome and, consequently, improved clearance of the pathogen. This information opens new avenues to target the immune response to *C. burnetii* infection with the goal to eliminate the bacteria and thereby prevent disease.

## INTRODUCTION

The zoonotic infection Q fever is caused by the obligate intracellular Gram-negative bacterium *Coxiella* (*C*.) *burnetii*. The main reservoir of *C. burnetii* are small ruminants, but other mammals, birds or arthropods are also infected (1). Q fever is a relatively rare human infection, but large outbreaks can occur, such as in the Netherlands between 2007 and 2011 with more than 4000 infected individuals (2). In humans, 40% of the infected patients exhibit flu-like symptoms, such as fever and headaches, but the acute infection can also result in pneumonia or hepatitis. Acute Q fever mainly resolves spontaneously or within a few weeks with antibiotic treatment (1, 3). Nevertheless, up to 5% of infected people may develop chronic Q fever, which usually manifests as endocarditis. Treatment of chronic Q fever requires at least 18 months of doxycycline and chloroquine administration (4). This prolonged treatment clearly shows the need to improve the knowledge base concerning *C. burnetii* pathogenesis to allow the development of novel treatment options.

The primary host cells of *C. burnetii* are alveolar macrophages. However, infections have also been detected in trophoblasts, endothelial and epithelial cells, and fibroblasts (5). Macrophages ingest the bacteria via phagocytosis into a phagosome. This vacuolar compartment matures by fusion with endosomes and lysosomes, forming a degradation-active phagolysosome, in which most bacteria are digested (6, 7). The *C. burnetii*-containing vacuole (CCV) matures along the canonical phagosome maturation process, with two exceptions: the CCV fuses with autophagosomes early after uptake and later with secretory vesicles (8, 9). Importantly, *C. burnetii* not only withstands the harsh environment of the phagolysosomal compartment, but actually requires it for the activation of its type IVB secretion system (T4BSS) (10). The T4BSS is an essential virulence factor, which mediates the translocation of bacterial effector proteins into the host cell cytosol (11). These effector proteins modulate several important host cell pathways for the benefit of the pathogen. As *C. burnetii* is a slowly replicating pathogen, it depends on host cell survival for maintaining its replicative niche (12). The underlying mechanisms required to maintain host cell viability are not yet completely understood. However, a functional T4BSS is essential for this activity, and several effector proteins modulating host cell survival have been described (13, 14). The effector proteins AnkG, CaeA and CaeB inhibit apoptosis (15–18), whereas the effector protein IcaA inhibits pyroptosis (19).

Here, we focus on the interplay of *C. burnetii* and pyroptosis. Pyroptosis is a pro-inflammatory form of cell death, which serves as an important mechanism to eliminate pathogens (20). There are two pathways leading to pyroptosis, the canonical and the non-canonical inflammasome pathways, which crosstalk during infection (21, 22). The canonical pathway is activated by recognition of damage-associated molecular patterns (DAMPs) or pathogen-associated molecular patterns (PAMPs) by inflammasome receptors. NLRP3 (NOD-, LRR- and pyrin domain-containing protein 3) is one of a variety of such receptors (21). Activation of these receptors leads to their oligomerization, which results in the recruitment of the adaptor protein ASC. This leads to the subsequent recruitment of pro-caspase 1, which is processed through autoproteolytic cleavage to yield the active caspase 1. Caspase 1 then cleaves pro-IL1β, pro-IL18 and gasdermin D (GSDMD) (23). The N-terminal part of cleaved GSDMD binds to phosphoinositides in the plasma membrane, where its oligomerization leads to the formation of pores (24). These pores allow the release of mature IL1β and disrupt the osmotic potential, which eventually results in cell lysis (25). In contrast, the non-canonical pathway is activated by intracellular lipopolysaccharides (LPS) from Gram-negative bacteria, which is recognized by GBP1, a member of the dynamin-related guanylate binding proteins (GBPs) (26). In mice, GBP1-bound LPS initiates the assembly of caspase 11 (27), while in humans, GBP2-bound LPS coordinates caspase 4 recruitment and activation (28). This results in oligomerization of the catalytic subunits of caspase 11 and acquisition of protease function, leading to cleavage of GSDMD and subsequent pore formation, which allows secretion of cytoplasmic molecules and potentially cell lysis (27, 29). In addition, the induction of the non-canonical inflammasome results in the activation of the canonical caspase 1-dependent inflammasome (30, 31). Whether the infection of macrophages with *C. burnetii* activates the inflammasome (32) or not (19, 33, 34) is controversial. Additionally, one study demonstrated that IcaA, a T4BSS effector protein, inhibits caspase 11 activation (19), while another study suggested that the T4BSS has no influence on inflammasome activation or inhibition (34). The aim of this study was, therefore, to investigate the interplay of *C. burnetii* and the non-canonical inflammasome, and how this is influenced by the T4BSS and oxygen availability.

## RESULTS

### *C. burnetii* induces IL1β secretion in a dose- and growth phase-dependent manner

First, it was evaluated whether *C. burnetii* induces secretion of IL1β by murine BMDM and whether its secretion level is affected by the growth phase or the multiplicity of infection (MOI). Therefore, the different growth phases (lag, log or stationary) for *C. burnetii* Nine Mile phase II (wt) and the corresponding T4BSS mutant (Δ*dotA*) in ACCM-2 media were defined (Fig. 1A). Subsequently, BMDM were infected with different MOIs of bacteria grown to the three growth phases. Notably, a dose-dependent increase in IL1β secretion was detected after infection with lag-and log-phase grown wt and Δ*dotA* bacteria, which was not the case for bacteria grown to the stationary phase (Fig. 1B). *C. burnetii* is highly infectious and less than 6 bacteria might cause illness in humans (35). Therefore, an MOI of three was used for the remainder of the study and the effect of the growth phase on the induction of IL1β secretion was compared (Fig. 1C). The infection with lag or log phase grown bacteria induced similar amounts of IL1β secretion, in contrast to stationary phase bacteria (Fig. 1C). Consequently, log phase grown *C. burnetii* was used in all further experiments.

**Figure 1:**
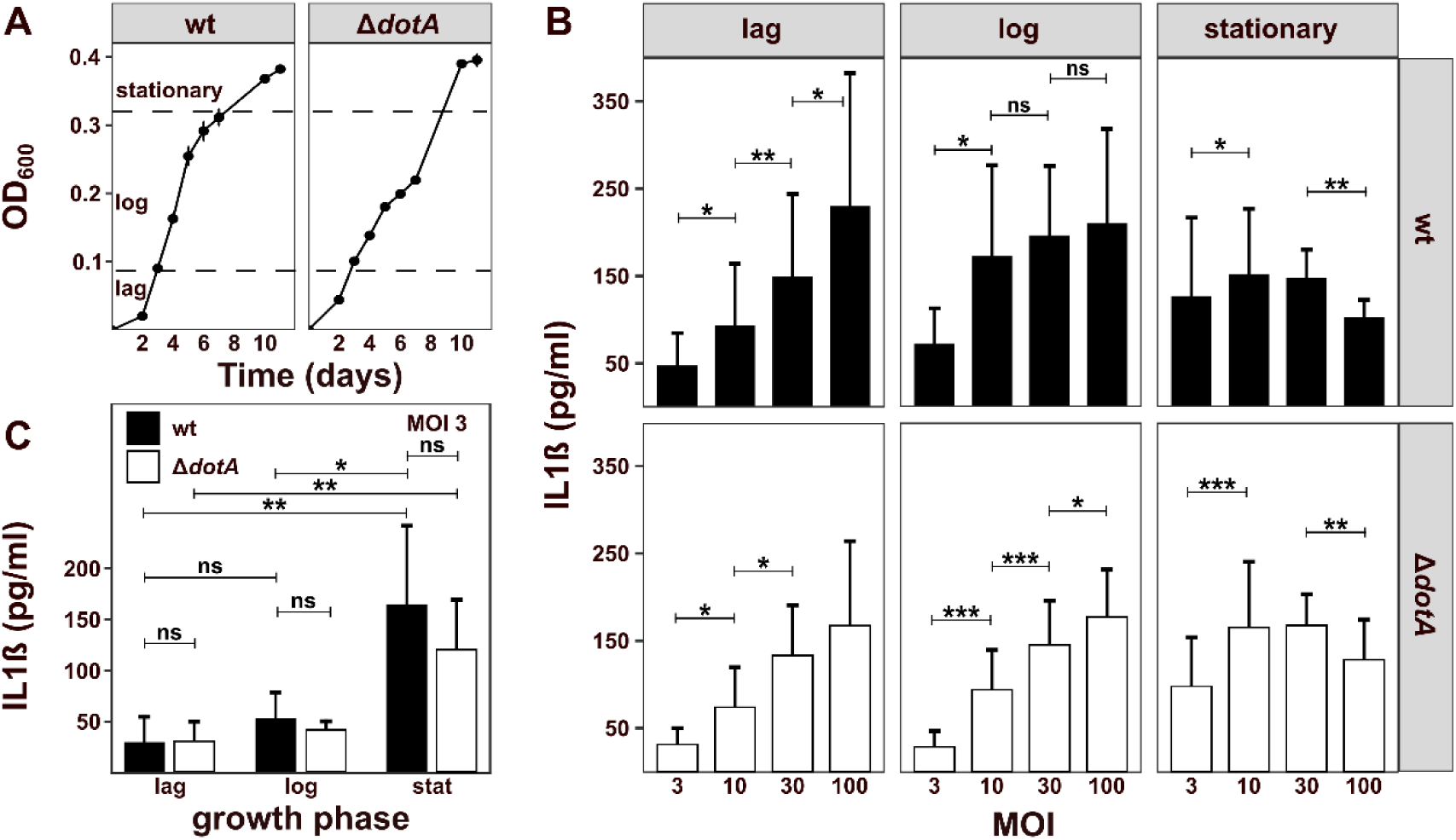
The induction of the inflammasome is dependent on the growth phase and MOI of *C. burnetii*. (A) The growth curves of the *C. burnetii* wt and Δ*dotA* strains were determined by culturing the bacteria for 12 days with regular measurement of the optical density at 600 nm (OD_600_). (B) BMDM were infected for 32h with MOIs of 3, 10, 30 or 100 of the *C. burnetii* wt and Δ*dotA* strains grown to lag, log, and stationary growth phases. Secretion of IL1β was quantified by ELISA. (C) IL1β secretion was examined in murine BMDM infected at MOI 3 with *C. burnetii* wt and Δ*dotA*, which had been cultivated to the different growth phases. Mean ± SD, n = 3, t-test. ns = non significant, *p < 0.05, **p < 0.01, ***p < 0.001.

### *C. burnetii* only slightly induces IL1β expression and secretion

In order to determine the strength of IL1β expression and secretion induced by *C. burnetii*, it was compared to the response induced by outer membrane vesicles from *Escherichia coli* (OMVs) and *L. pneumophila* Δ*flaA*, both known activators of the non-canonical inflammasome pathway. To evaluate whether *C. burnetii* is able to inhibit the non-canonical inflammasome in the presence of potent inducers, *C. burnetii* infected cells were treated with OMVs or *L. pneumophila* Δ*flaA* (36, 37). The data demonstrate that *C. burnetii* induced only minor levels of both IL1β mRNA (Fig. 2A) and secreted IL1β protein (Fig. 2B), which is in line with previous reports (19, 34). Induction of IL1β expression and secretion by *C. burnetii* was independent of the presence of a functional T4BSS (Fig. 2A and B). Evidently, the prior infection with wt or Δ*dotA C. burnetii* augmented OMV- and *L. pneumophila* Δ*flaA*-induced IL1β expression and secretion (Fig. 2A and B). This finding is consistent with the results of Delaney *et al*., 2021, but contrasts Cunha *et al*., 2015. Taken together *C. burnetii* only slightly induces IL1β expression and secretion, but may boost IL1ß levels by other stimuli, independent of its T4BSS. These data suggest that *C. burnetii* is unable to inhibit the non-canonical inflammasome.

**Figure 2:**
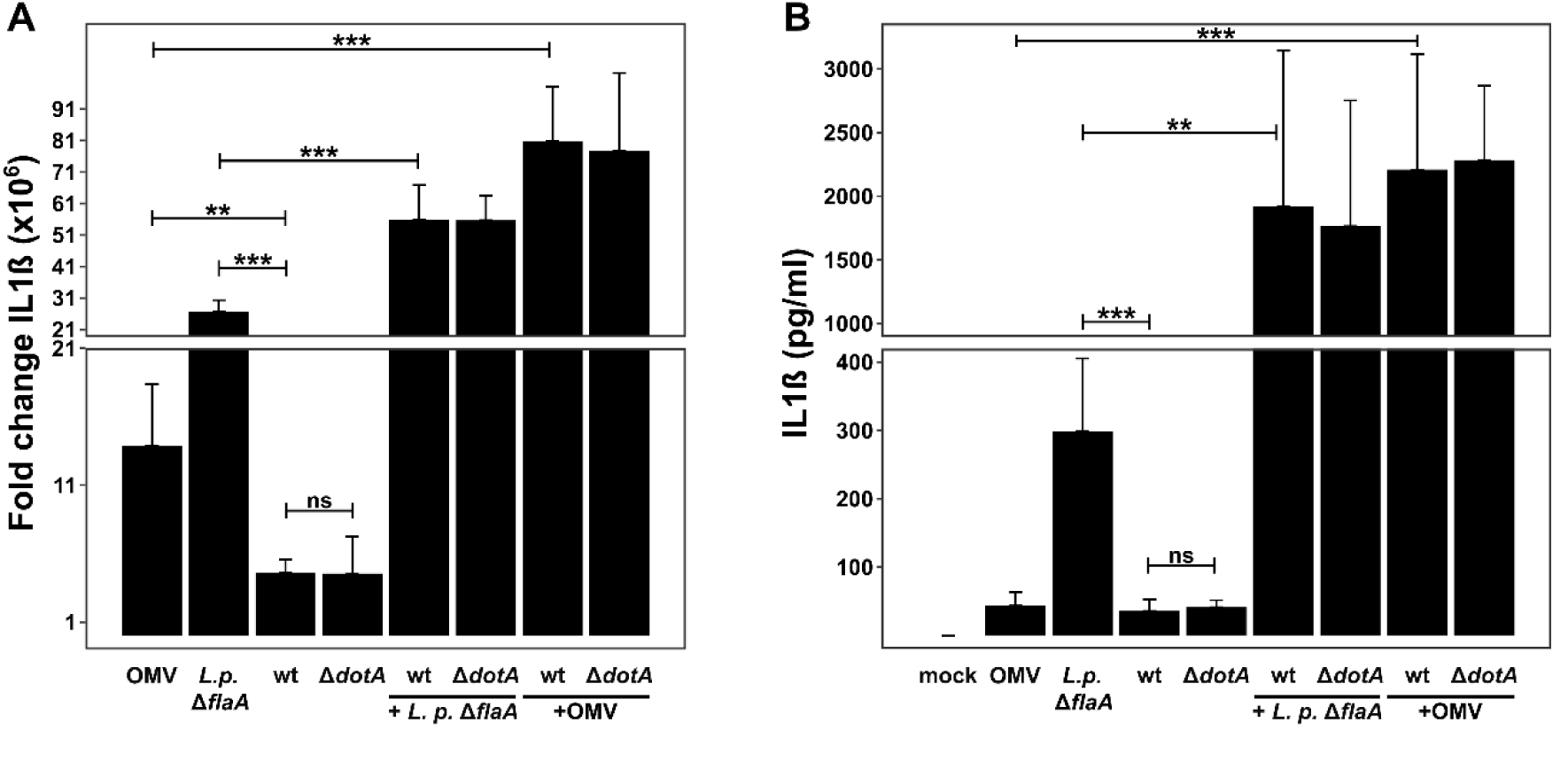
*C. burnetii* fails to potently induce IL1ß transcription and secretion. BMDM were stimulated with OMVs (2 µg/ml) for 8h, infected with *L. pneumophila* Δ*flaA* for 8h, infected with *C. burnetii* wt or Δ*dotA* for 32h or infected with *C. burnetii* wt or Δ*dotA* for 24h with subsequent infection with *L. pneumophila* Δ*flaA* or stimulation with OMVs for 8h. (A) The fold change of the IL1β mRNA level relative to uninfected BMDM was determined by RT-qPCR analysis. Mean ± SD, n = 3, t-test. (B) Secretion of IL1β was quantified by ELISA. Mean ± SD, n = 3, Mann-Whitney U test. **p < 0.01, ***p < 0.001.

### *C. burnetii* does not inhibit OMV or Δ*flaA*-induced activation of the non-canonical inflammasome

To test whether *C. burnetii* is unable to inhibit the non-canonical inflammasome, the non-canonical and the canonical inflammasome pathways were analyzed by immunoblot analysis, as activation of the non-canonical inflammasome results in activation of the canonical inflammasome pathway. Therefore, BMDM were infected with wt or Δ*dotA C. burnetii* alone or in combination with OMVs or *L. pneumophila* Δ*flaA*. Cell lysates were analyzed for the expression of GBP1, caspase 11, cleaved GSDMD, NLRP3, IL1β (uncleaved and cleaved) with actin serving as loading control. The infection with *C. burnetii* resulted in induced expression of GBP1, caspase 11 and NLRP3, and lack of cleaved GSDMD (Fig. 3), suggesting that *C. burnetii* induced the non-canonical inflammasome only insufficiently. This was supported by reduced oligomerization of caspase 11 in *C. burnetii* infected cells in comparison to cells infected with *C. burnetii* in combination with *L. pneumophila* Δ*flaA* (Fig. S1). Furthermore, expression of uncleaved IL1β was not detectible, suggesting reduced transcription of IL1ß. Interestingly, *C. burnetii* boosted the activation of the non-canonical inflammasome by an additional stimulus (OMV or *L. pneumophila* Δ*flaA*), in a T4BSS-independent manner (Fig. 3).

**Figure 3:**
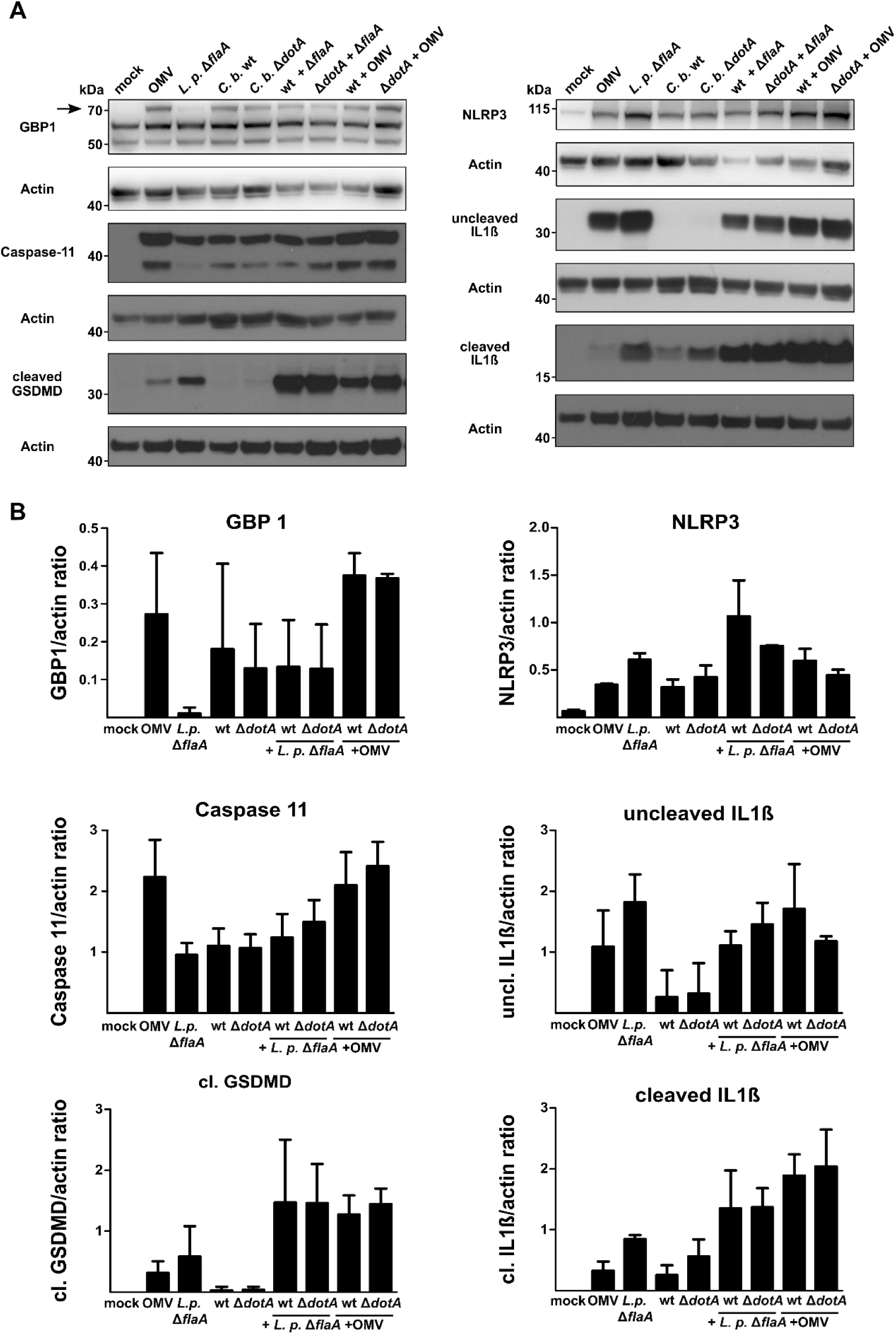
Lack of potent non-canonical inflammasome activation by *C. burnetii* can be overcome by additional stimulation. BMDM were stimulated with OMVs (2 µg/ml) for 8h, infected with *L. pneumophila* Δ*flaA* for 8h, infected with *C. burnetii* wt or Δ*dotA* for 32h or infected with *C. burnetii* wt or Δ*dotA* for 24h with subsequent infection with *L. pneumophila* Δ*flaA* or stimulation with OMVs for 8h. Immunoblot analysis was performed using antibodies against GBP1, caspase 11, cleaved GSDMD, NLRP3, uncleaved IL1β, cleaved IL1β, and actin as loading control. One representative blot from at least four independent experiments is shown for each target.

### Hypoxia hampers inflammasome activation by different stimuli

We previously demonstrated that a *C. burnetii* infection under low levels of oxygen (hypoxia) leads to transcriptional upregulation of pro-inflammatory cytokines (38). As bacterial replication was observed under hypoxia (39), we hypothesized that hypoxia might induce inflammasome activation, contributing to the control of *C. burnetii*. This hypothesis is supported by the observation that the infection of macrophages with *Helicobacter pylori* resulted in enhanced caspase 1 activation and IL1β secretion under hypoxia (40). To test whether hypoxia might induce inflammasome activation, IL1β-transcription and -secretion were determined under hypoxic (0.5% O_2_) conditions. Similar to the results obtained under normoxic conditions (21% O_2_) (Fig. 2A), *C. burnetii* induced only low levels of IL1β mRNA and the prior infection with wt and Δ*dotA C. burnetii* augmented OMV- and *L. pneumophila* Δ*flaA*-induced IL1β expression (Fig. 4A). However, to determine the impact of oxygen availability, the fold change in IL1β mRNA levels induced by the various stimuli under normoxia (Fig. 2A) and hypoxia (Fig. 4A) were compared. The data revealed a strong reduction of IL1β transcription under hypoxic conditions (Fig. 4B), suggesting a general block of IL1β transcription induction by different stimuli of the non-canonical inflammasome under hypoxia. This block was also translated to the amount of secreted IL1β. Accordingly, only low levels of secreted IL1β were detected by doubly stimulated macrophages (OMV + *C. burnetii*) under hypoxia. Thus, doubly stimulated macrophages produced ∼40pg/ml under hypoxia (Fig. 4C) versus ∼2000pg/ml under normoxia (Fig. 2B). These data indicate that hypoxia might restrict the inflammasome response of macrophages. To address this experimentally, the non-canonical inflammasome pathway was analyzed by immunoblot analysis. While caspase 11 expression was induced, cleavage of GSDMD was not detected in any of the samples analyzed (Fig. 5). Furthermore, uncleaved IL1β was detected, but the cleaved form of IL1ß was absent in cells treated with a single stimulus (Fig. 5). These results suggest that hypoxia blocks activation of the non-canonical inflammasome. Nonetheless, it does not address the question why *C. burnetii* induces the non-canonical inflammasome only weakly.

**Figure 4:**
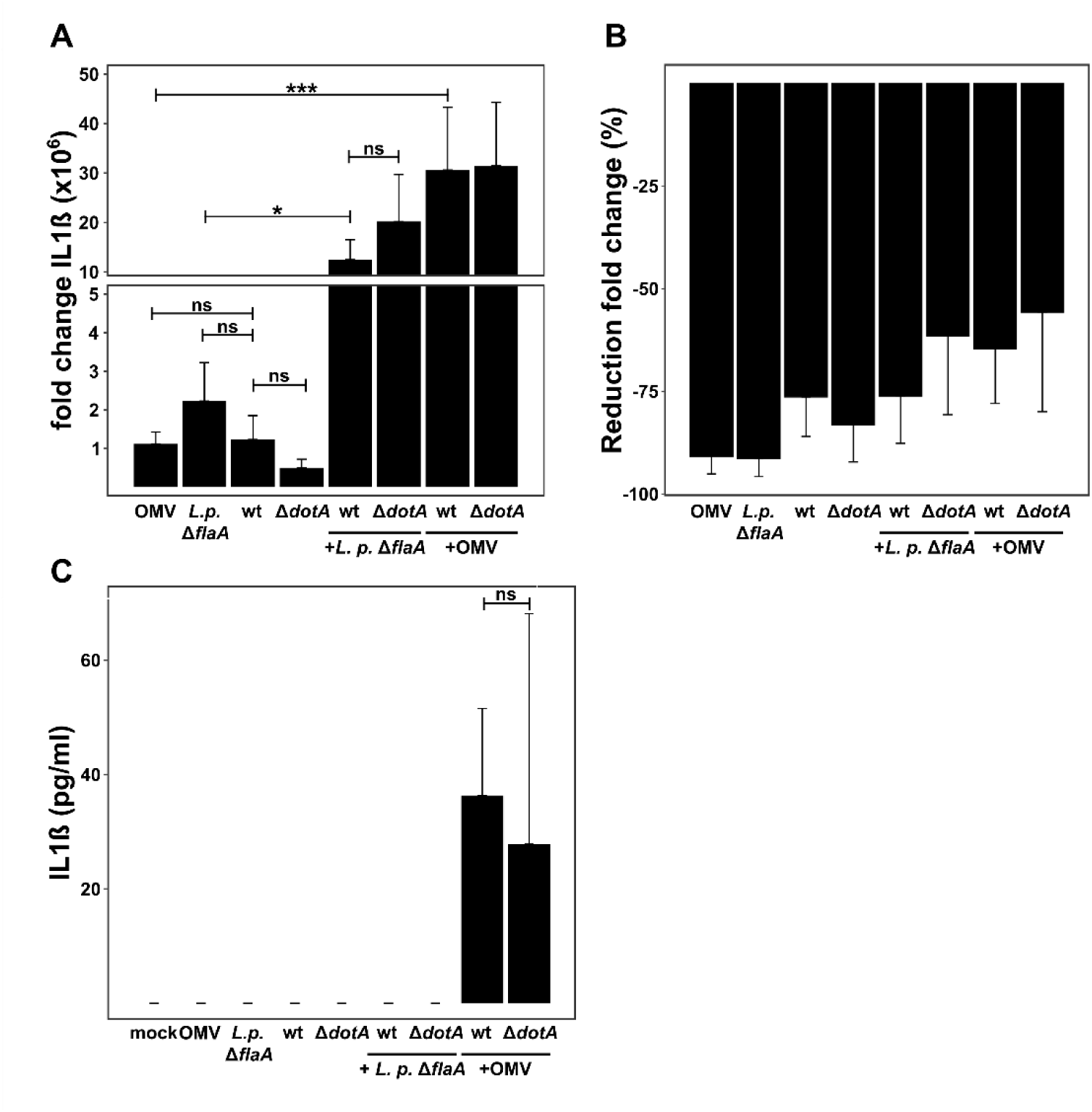
Transcription and secretion of IL1β is reduced under hypoxic conditions. BMDM were stimulated with OMVs (2 µg/ml) for 8h, infected with *L. pneumophila* Δ*flaA* for 8h, infected with *C. burnetii* wt or Δ*dotA* for 32h or infected with *C. burnetii* wt or Δ*dotA* for 24h with subsequent infection with *L. pneumophila* Δ*flaA* or stimulation with OMVs for 8h under hypoxic conditions (0.5% O_2_). (A) Fold change of the IL1β mRNA transcript levels relative to uninfected BMDM were quantified by RT-qPCR. Mean ± SD, n = 3, One-Way-ANOVA with Bonferroni post-test. (B) Reduction rate of the IL1β mRNA transcript level under hypoxic condition relative to the infection under normoxic (21% O_2_) conditions. (C) IL1β secretion was determined by ELISA. Mean ± SD, n = 3, Mann-Whitney U test. *p < 0.05, ***p < 0.001.

**Figure 5:**
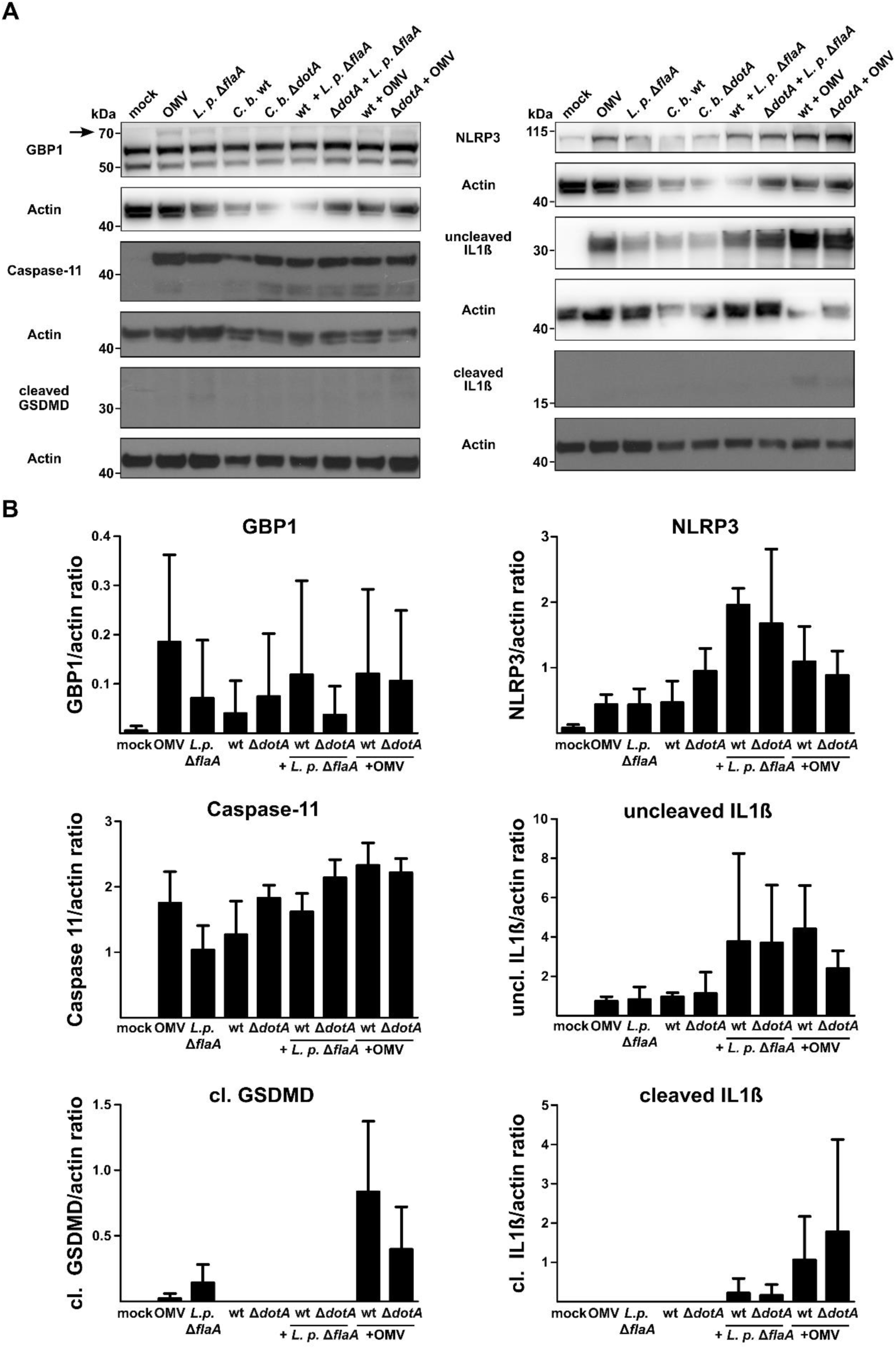
Inflammasome activation is strongly reduced under hypoxic conditions. BMDM were stimulated with OMVs (2 µg/ml) for 8h, infected with *L. pneumophila* Δ*flaA* for 8h, infected with *C. burnetii* wt or Δ*dotA* for 32h or infected with *C. burnetii* wt or Δ*dotA* for 24h with subsequent infection with *L. pneumophila* Δ*flaA* or stimulation with OMVs for 8h under hypoxic conditions (0.5% O_2_). Immunoblot analysis was performed using antibodies against GBP1, caspase 11, cleaved GSDMD, NLRP3, uncleaved IL1β, cleaved IL1β, and actin as loading control. One representative blot from at least four independent experiments is shown for each target. Importantly, the same membrane was used to analyze the expression of GBP1 and NLRP3. Therefore, the actin blot is the same.

### TRIM21 is only slightly upregulated by infection with *C. burnetii*

As the infection with *C. burnetii* induced expression of caspase 11, but not of cleaved GSDMD, possible mechanism(s) underlying this observation were investigated. Therefore, the transcriptional levels of possible modulators of caspase 11 activity or GSDMD cleavage were determined after infection with *C. burnetii*, treatment with OMV or a combination of both. We focused on the following proteins: Gate-16, Irgm2, Serpinb1 and TRIM21. The ATG8 family member Gate-16 and the IFN-inducible protein Irgm2 cooperatively restrict the activation of the non-canonical inflammasome in macrophages by inhibiting caspase 11-mediated cytokine release and pyroptosis (41, 42). The protease inhibitor Serpinb1 inhibits cathepsin G, which is involved in GSDMD cleavage (43). In contrast to Irgm2, Gate-16 and Serpinb1, which all inhibit non-canonical inflammasome activation, the tripartite motif protein TRIM21 acts as a positive regulator of GSDMD-mediated pyroptosis (44). As shown in figure 6, the expression of Gate-16 was not altered by any treatment. However, OMV treatment with or without prior infection with *C. burnetii* resulted in increased expression of Irgm2 and TRIM21, but did not alter the expression level of Serpinb1. Regardless of the presence of a functional T4BSS, *C. burnetii* infection resulted in a minor upregulation of TRIM21 and Irgm2 expression, but inhibited the expression of Serpinb1. These results demonstrate that *C. burnetii* does not interfere with OMV-induced changes in the expression levels of the genes analyzed. Moreover, the limited induction of TRIM21 expression might contribute to the inefficient activation of the non-canonical inflammasome by *C. burnetii*.

**Figure 6:**
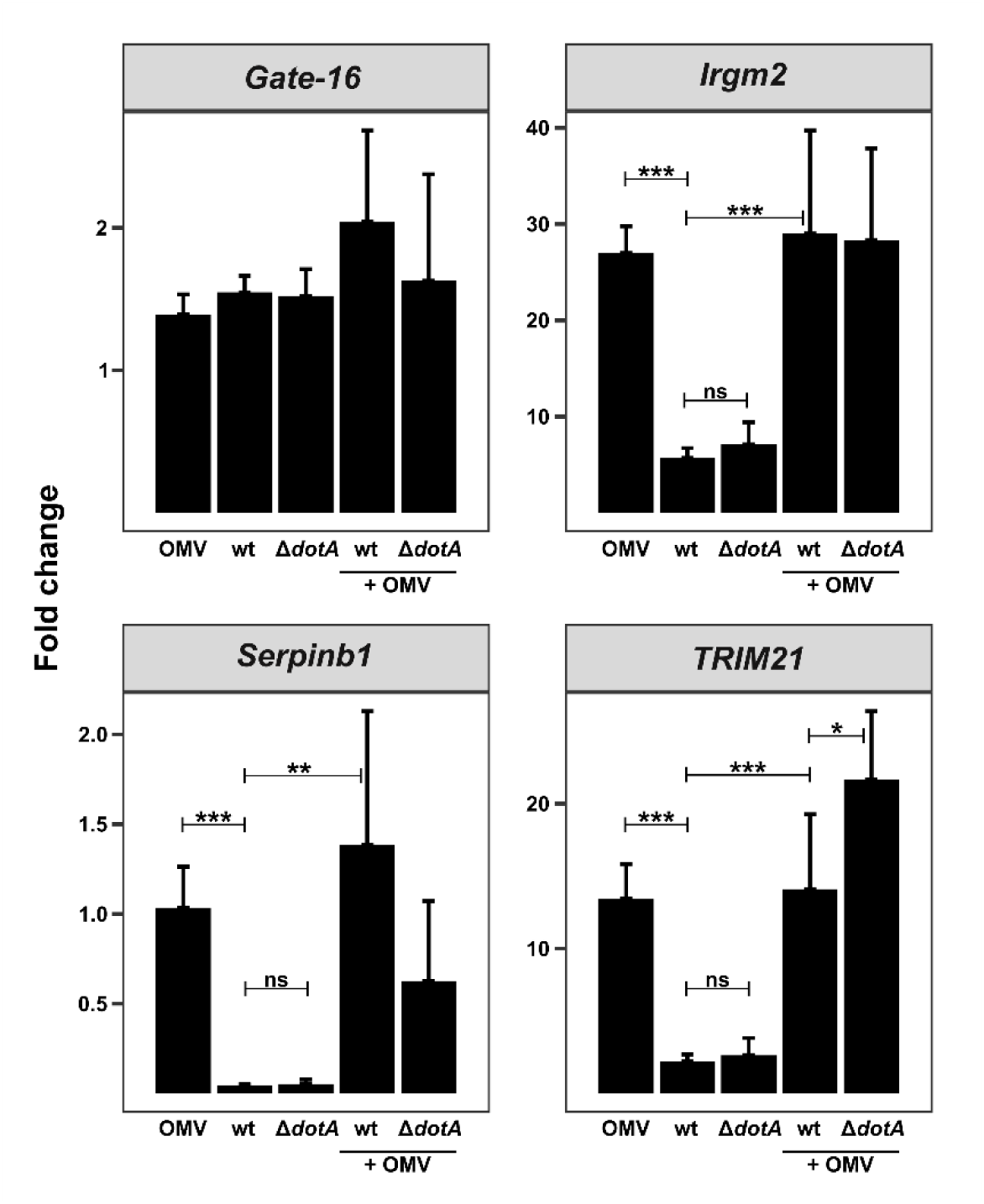
TRIM21 might be implicated in the lack of non-canonical inflammasome activation by *C. burnetii*, unlike GATE-16, Irgm1-3, or SerpinB1. BMDM were stimulated with OMVs for 8h, infected with *C. burnetii* wt or ΔdotA for 32h, or for 24h with subsequent OMV stimulation for 8h. The fold change in mRNA transcript level of the indicated targets was calculated relative to uninfected BMDM. Mean ± SD, n = 4, One-Way-ANOVA with Bonferroni post-test. **p < 0.01, ***p < 0.001.

### Itaconate does not mediate the inefficient activation of the non-canonical inflammasome by C. burnetii

The metabolite itaconate was reported to inhibit the NLRP3 inflammasome by preventing GSDMD processing (45, 46). However, contrary results have also been published (47). Itaconate is synthesized via decarboxylation of cis-aconitate by the enzyme immune-responsive gene 1 (IRG1), also termed ACOD1 (48). Itaconate was first identified as anti-microbial, but later also as immunoregulatory (48–50). Importantly, we previously demonstrated that a *C. burnetii* infection of murine macrophages induces the production of itaconate (39). Based on these findings, we hypothesized that itaconate might contribute to the inefficient activation of the non-canonical inflammasome by *C. burnetii*. Thus, IL1β secretion by *Acod1*^-/-^ and control BMDM was analyzed. Treatment with OMVs after prior infection with *C. burnetii* resulted in measurable levels of IL1β, while LPS + nigericin treatment resulted in a stronger IL1β response. However, the *C. burnetii* infection did not induce increased IL1β secretion by *Acod1*^-/-^ BMDM. Instead, secretion of IL1β was reduced in *Acod1*^-/-^ BMDM under all conditions tested (Fig. 7A), suggesting that itaconate might not inhibit activation of the non-canonical inflammasome. This hypothesis was further supported by the absence of any differences in caspase 11 expression or GSDMD cleavage between control and *Acod1*^-/-^BMDM (Fig. 7B). In contrast, a difference in the ability of *C. burnetii* to replicate in control versus *Acod1*^-/-^ BMDM was observed (Fig. 7C). As previously shown, the lack of itaconate allowed the replication of wt *C. burnetii* in *Acod1*^-/-^ BMDM, but not of the Δ*dotA* mutant (51). Importantly, addition of OMVs to wt or Δ*dotA C. burnetii* infected *Acod1*^-/-^ BMDM reduced the bacterial load (Fig. 7C). These data suggest that activation of the non-canonical inflammasome may help to control *C. burnetii*.

**Figure 7:**
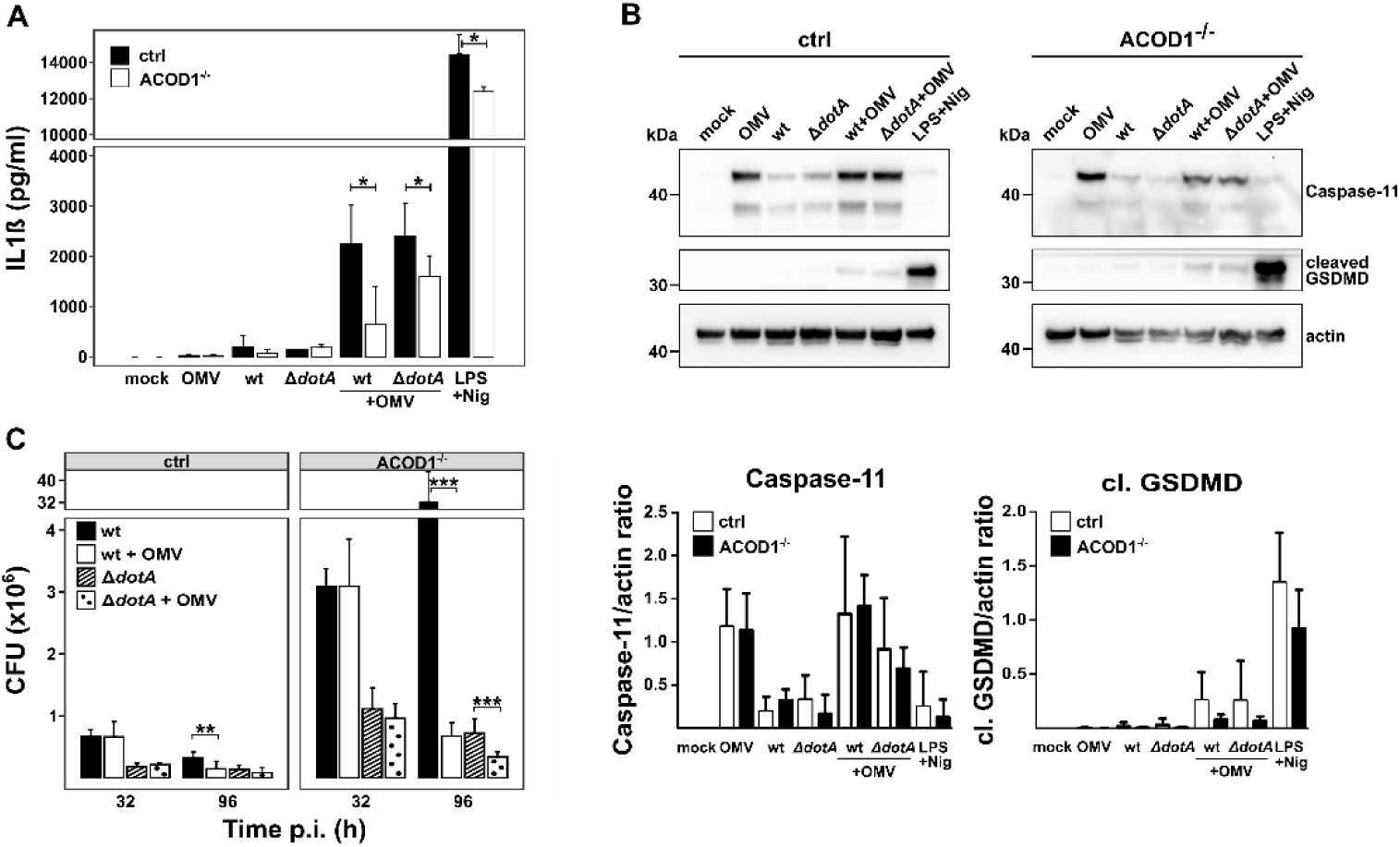
Itaconate deficiency facilitates extensive *C. burnetii* replication, but does not enable increased inflammasome activation. (A-B) ACOD1^+/-^ (ctrl) and ACOD1^-/-^ BMDM were stimulated with OMVs, LPS + Nigericin, or infected with wt and Δ*dotA C. burnetii* at MOI 3 with or without additional stimulation with OMVs. IL1ß secretion was quantified by ELISA (mean ± SD, n = 3, Mann-Whitney U test). Immunoblot analysis was conducted using antibodies against caspase 11, cleaved GSDMD, and actin as loading control. One representative blot of three independent experiments is shown. (C) Infection of ACOD1^+/-^ (ctrl) and ACOD1^-/-^ BMDMs with wt and Δ*dotA C. burnetii* were either not treated or stimulated with OMV. CFU count was determined at the time points indicated. Mean ± SD, n = 3, t-test. *p < 0.05, **p < 0.01, ***p < 0.001.

### Activation of the non-canonical inflammasome in *C. burnetii* infected BMDM resulted in induction of pyroptosis and bacterial clearance

To determine how OMV-induced inflammasome activation of *C. burnetii* infected BMDM led to bacterial clearance, the induction of pyroptosis was analyzed. Indeed, OMV-treatment of *C. burnetii*-infected BMDM resulted in decreased cell viability (Fig. 8A) and increased LDH release (Fig. 8B). This data suggests that OMV-mediated activation of the non-canonical inflammasome of *C. burnetii*-infected BMDMs resulted in pyroptosis and had significant impact on the ability of *C. burnetii* to survive. Indeed, reduced bacterial numbers were detected in both infected cells (Fig. 9A) and supernatants (Fig. 9B). These results demonstrate that efficient activation of the non-canonical inflammasome allowed the control of *C. burnetii*. Hence, the ability of *C. burnetii* to eschew non-canonical inflammasome activation represents an important immune-evasion strategy of this obligate intracellular pathogen.

**Figure 8:**
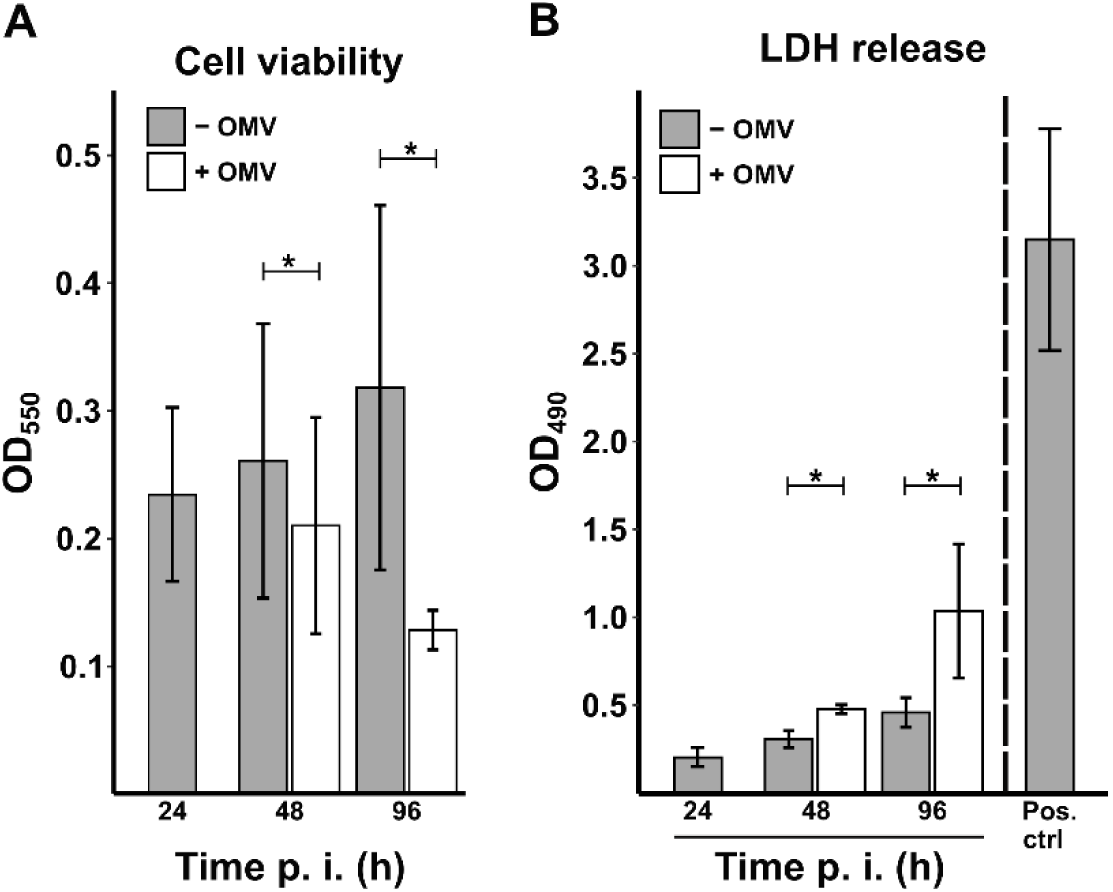
Activation of the inflammasome leads to induction of cell death. BMDM were infected with *C. burnetii* at MOI 3. 24h post infection, the cells were either not-treated or treated with 2 µg/ml OMVs. (A) At 24, 48 and 96h post-infection, the supernatant was removed and cell viability assessed using a crystal violet assay. (B) Supernatant was collected at 24, 48 and 96h post infection and an LDH assay was conducted determining LDH release. Mean ± SD, n = 3 - 4, each as duplicates or triplicates, Mann-Whitney U test. *p < 0.05, ***p < 0.001.

**Figure 9:**
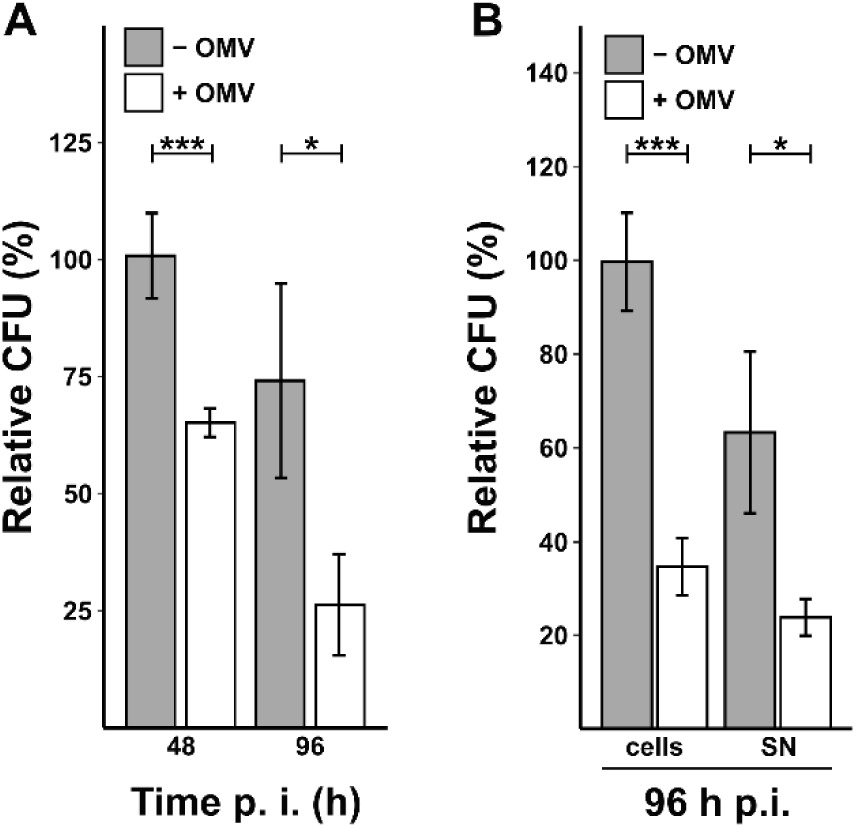
Inflammasome activation results in a reduced bacterial burden. BMDM were infected with *C. burnetii* at MOI 3. 24h post-infection, the cells were either left untreated or treated with 2 µg/ml OMVs. (A) Bacteria were isolated from the cells 48 and 96h post-infection, and their CFU count relative to the 48h untreated sample was determined. (B) Bacteria were isolated 96h post-infection from both cells and supernatant (SN). The CFU count was calculated relative to the CFU count determined from the untreated cells. Mean ± SD, n = 3 – 4, each as duplicates or triplicates, t-test. *p < 0.05, ***p < 0.001.

### Acylation pattern of lipid A is decisive for evasion of inflammasome activation

We hypothesized that the immune-evasion strategy might be dependent on the chemical structure of lipid A, which is only tetra-acylated in *C. burnetii* (52). *C. burnetii* encodes all enzymes necessary for lipid A biosynthesis, except for LpxM and LpxL, which are required for adding the secondary acyl chains to the tetra-acylated lipid A. Activation of the non-canonical inflammasome requires penta- and hexa-acylated lipid A, while tetra-acylated lipid A acts as an antagonist and does not lead to activation (53, 54). To test the role of lipid A acylation, *C. burnetii* lipid A was modified by heterologous expression of the *Escherichia coli lpxL* and *lpxM* genes. This attempt was unsuccessful, possibly because acceptor molecules were missing. In order to provide appropriate acyl chains as acceptor molecules, all four acyltransferase encoding genes in *E. coli* - *lpxL*, *lpxM*, *lpxD* and *lpxA* - were expressed with either Flag-, HA- or His-tags under an inducible promotor (55). First, the expression of all four genes was verified by immunoblot using tag-specific antibodies (Fig. 10A). Next, the level of IL1ß secretion induced by OMV, infection with wild-type *C. burnetii* or two different mutants, expressing *E. coli* LpxA, LpxD, LpxM and LpxL was determined. The two mutants induced an increased amount of IL1ß compared to wild-type *C. burnetii* (Fig. 10B). Notably, the two mutants also showed reduced survival, which was not further reduced by addition of OMVs (Fig. 10C). These data suggest that the expression of *E. coli* LpxA, LpxD, LpxM and LpxL in *C. burnetii* might change the acylation pattern of lipid A, allowing activation of the non-canonical inflammasome, and consequently promoting a better control of infection. However, how this improved activation of the non-canonical inflammasome results in *C. burnetii* control, has to be clarified.

**Figure 10:**
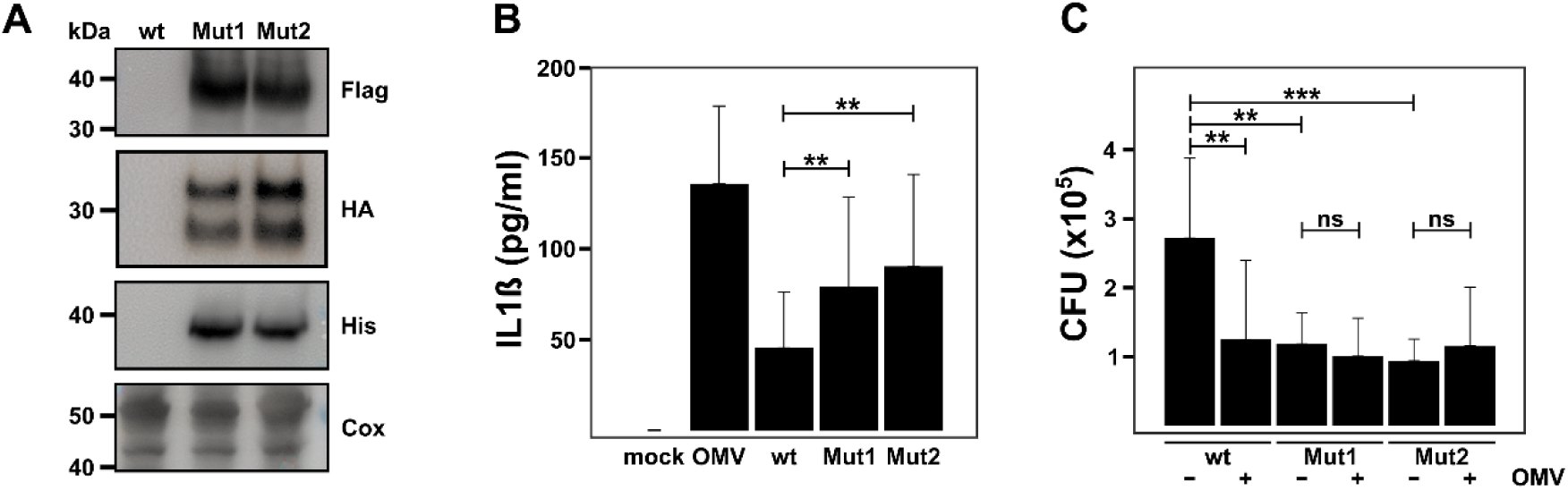
Altered acylation pattern of lipid A results in activation of the inflammasome. (A) with *C. burnetii* wild-type (WT) or two different mutants expressing LpxA, LpxD, LpxL and LpxM (Mut1 and Mut2) were subjected to immunoblot analysis using antibodies against Flag (to detect LpxD (∼39 kDa)), HA (to detect LpxA (∼28 kDa) and LpxL (∼36 kDa)), His (to detect LpxM (∼38 kDa) and *C. burnetii*. (B-C) BMDM were infected with *C. burnetii* wild-type (WT) or two mutants expressing LpxA, LpxD, LpxL and LpxM at a MOI of 3 for 32h. As controls, BMDM were either not treated (-) or treated for 8h with 2 µg/ml OMVs (+). (B) IL1ß secretion was quantified by ELISA (mean ± SD, n = 3, Wilcoxon signed rank test). (C) Bacteria were isolated from the cells 96h post-infection, and the CFU count was determined (mean ± SD, n = 3, each as triplicates, Mann-Whitney U test). **p < 0.01, ***p < 0.001).

To determine the lipid A acylation pattern of the mutant, MALDI-MS/MS analyses in negative ion mode using a Norharmane matrix was performed (Fig. 11 and Tab. S1). The molecular ion at m/z 1811.301 was identified as a hexaacylated lipid A composed of one 3-hydroxyhexadecanoic acid (3-OH-C16:0), one 3-hydroxypentadecanoic acid (3-OH-C15:0), two 3-hydroxytetradecanoic acid (3-OH-C14:0) and two dodecanoic acid (C12:0) chains (Fig. 12A). MS/MS fragmentation revealed characteristic low-mass fragment ions at m/z 198.4, 242.6, 255.7, and 269.7, consistent with the presence of free fatty acids C12:0, 3-hydroxy-C14:0, 3-hydroxy-C15:0 and 3-hydroxy-C16:0, respectively. The molecular ion detected at m/z 1628.508 was assigned to a pentaacylated lipid A species, comprising one 3-OH-C16:0, one 3-OH-C15:0, two 3-OH-C14:0 and one C12:0 fatty acid chain (Fig. 12B). The ions observed at m/z 1366.086 and 1446.123 correspond to tetraacylated lipid A molecules containing one and two phosphate groups, respectively. Both of these tetraacylated species share an acylation pattern consisting of one 3-OH-C16:0, one 3-OH-C15:0 and two 3-OH-C14:0 acid chains (Fig. 12C). These results support the hypothesis of a heterogeneous clone population, in which the tetraacylated form is predominant, while hyperacylated species are present at minor levels.

**Figure 11:**
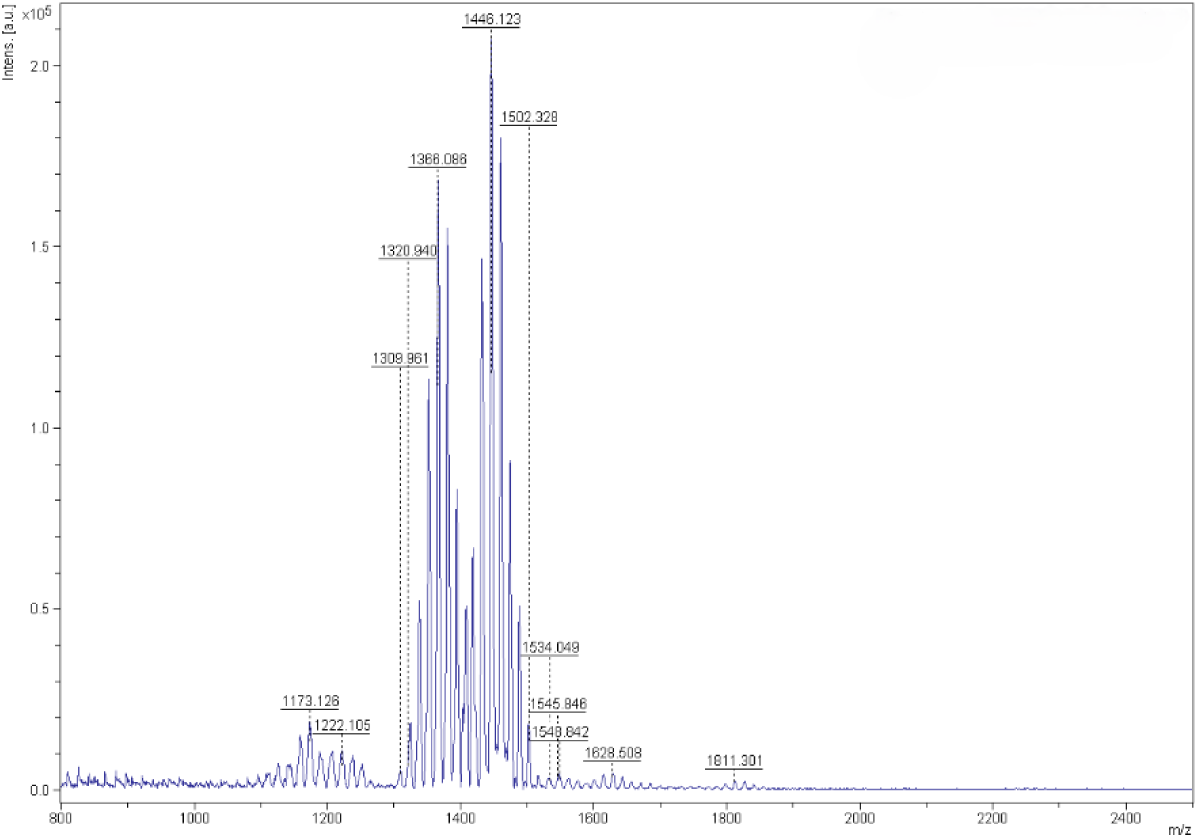
MALDI-MS spectrum of lipid A of *C. burnetii* mutant clone Mut1. Five liters of *C. burnetii* culture induced for 72h with 0.5 mM IPTG were fixed in 2% formaldehyde. LPS was isolated and applied to acetic acid hydrolysis. MALDI-TOF analyses were performed. Shown is one of two independent experiments.

**Figure 12:**
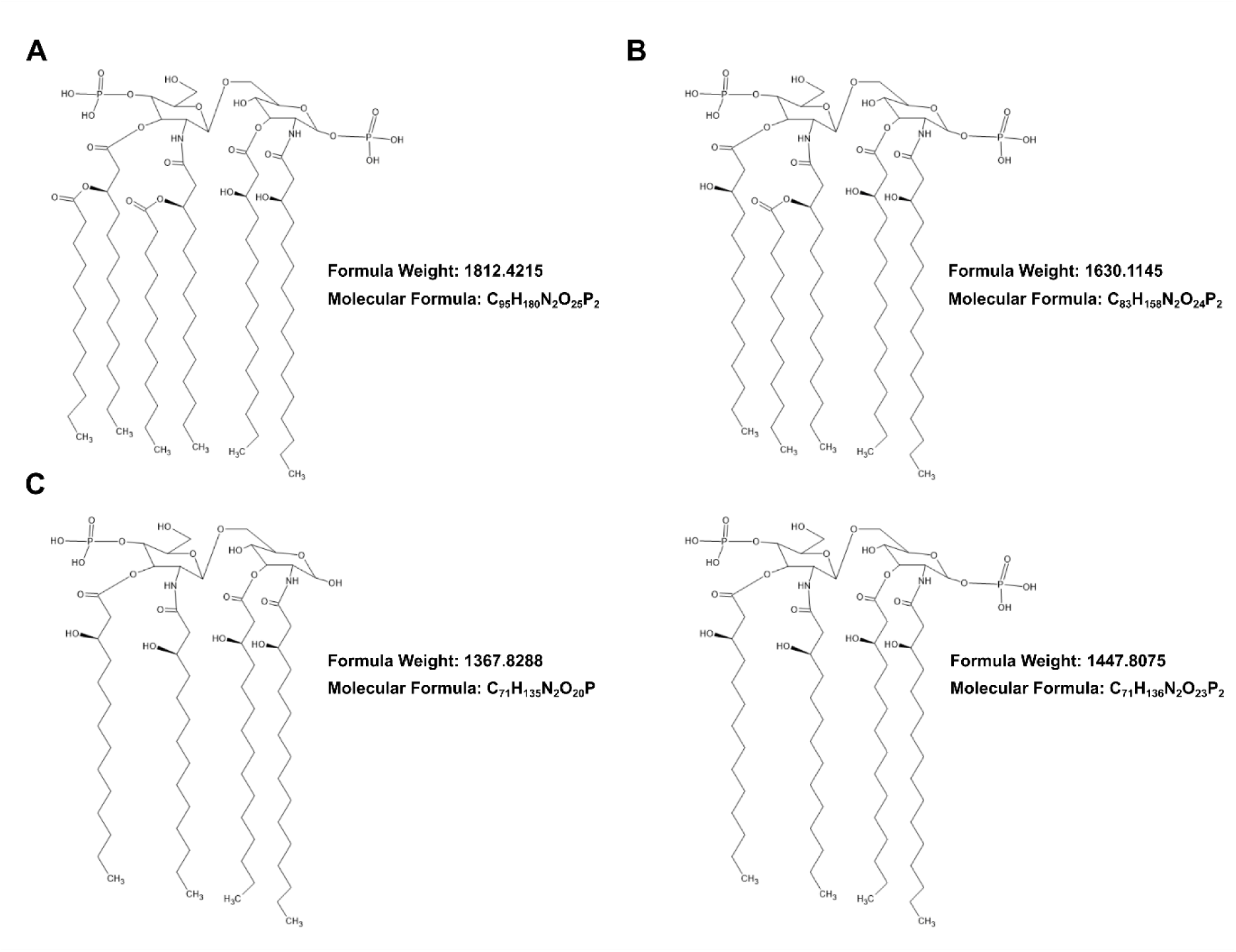
Chemical structure of the lipid A moiety of *C. burnetii* mutant clone Mut1 determined by MALDI-MS/MS analysis. (A) Hexaacylated lipid A. (B) Pentaacylated lipid A. (C) Tetraacylated lipid A, left: monophosphorylated lipid A, right: dephosphorylated lipid A.

## DISCUSSION

For *C. burnetii*, an obligate intracellular and slow-replicating pathogen, it is of utmost importance to manipulate host cell death pathways (14). To do so, *C. burnetii* uses different strategies (56). Thus, *C. burnetii* inhibits intrinsic and extrinsic apoptosis (57, 58) and T4BSS effector proteins are implicated in this process (15, 18, 59, 60). During late stages of a *C. burnetii* infection, apoptosis might be induced (61), which mediates bacterial egress (62). A *C. burnetii* infection also results in modulation of the non-canonical inflammasome, yet this phenomenon has not been studied as extensively as apoptosis modulation (56). The non-canonical inflammasome is a key host defense mechanism against Gram-negative bacteria, and therefore, it is necessary to investigate how *C. burnetii* evades this host immune response.

A bacterial infection might lead to reduced oxygen availability in the tissue (63), which in turn limits antimicrobial effector pathways of the host (64). The results obtained here suggest that inflammasome activation is also influenced by oxygen availability (Figs. 4 and 5). Severe hypoxia might prevent inflammasome activation and efficient clearance of the infection. Importantly, hypoxia can either promote or inhibit inflammasome-triggered pyroptosis. For example, hypoxia-mediated stabilization of HIF1α can lead to upregulation of caspase 1 and potentially, enhanced activation of the canonical inflammasome pathway. Conversely, HIF1α can upregulate PDK1 and thereby inhibit inflammasome activation (65). The cellular context is a decisive factor to determine the effect of hypoxia on inflammasome activation. In addition, bacterial virulence factors might influence hypoxia-mediated signaling, including HIF1α stabilization (66). Hence, *C. burnetii* dampens HIF1α stabilization in a T4BSS-dependent manner (38). Further research is required to determine how hypoxia prevents activation of the non-canonical inflammasome during *C. burnetii* infection.

The available information about the *C. burnetii* – non-canonical inflammasome interplay suggests that it might be strain-, host species- and/or even cell type-specific. For instance, in human alveolar macrophages, the virulent strain *C. burnetii* Nine Mile phase I (NM I) does not induce inflammasome activation, while the attenuated Nine Mile phase II (NM II) strain does; however, without inducing cell death (32, 67). This suggests, that the host response can differ depending on the bacterial strain. In addition, both host species and cell type might influence the host response to infection. NM II activates the inflammasome in human alveolar macrophages (32, 67) and murine B1a cells (68), but not in murine BMDM (19, 33, 34). The results presented in this study support the observation that NM II *C. burnetii* does not activate the non-canonical inflammasome in murine BMDM. Notably, while caspase 11 expression was detected during *C. burnetii* infection, the enzyme appears to be in an inactive state, as caspase 11 oligomerization was only partially induced (Fig. S1) and GSDMD cleavage was not observed (Figs. 2A, 2B, 3).

It has been controversially discussed whether *C. burnetii* actively inhibits non-canonical inflammasome activation or not, and whether the T4BSS is involved in this activity (19, 34). The data presented here indicate that *C. burnetii* does not activate the non-canonical inflammasome regardless of the presence or absence of the T4BSS, suggesting a passive rather than an active mechanism. Accordingly, *C. burnetii* is unable to inhibit activation of the non-canonical inflammasome induced by other stimuli, like OMVs or *L. pneumophila* Δ*flaA* (Figs. 2 and 3). These findings are consistent with a model of avoidance of, but not active interference with activation of the non-canonical inflammasome by *C. burnetii*. Several mechanisms utilized by pathogens to prevent inflammasome activation have been described. These mechanisms are partially dependent on virulence factors. In *Mycobacterium tuberculosis*, the serine threonine kinase PnkF inhibits inflammasome activation by inhibiting K^+^ and Cl^-^ efflux and reactive oxygen species production (69), while the *M. tuberculosis* hydrolase Hip1 dampens TLR2-dependent recognition and signaling, and thereby inflammasome activation (70). Further examples of microbial virulence factors, that actively interfere with inflammasome activation, have been described (71–74). In contrast, the results suggest that *C. burnetii* appears to avoid inflammasome activation, but is unable to prevent other stimuli from activating the non-canonical inflammasome pathway (Figs. 2 and 3), despite activation of this pathway resulting in improved control of *C. burnetii* (Fig. 9). The data also suggest that lipid A may contribute to this lack of efficient activation of the non-canonical inflammasome (Fig. 10). Lipid A, as a component of LPS, is the primary inducer of caspases 4, 5 and 11 (75, 76). The acylation pattern of lipid A determines its recognition by the immune system and non-canonical inflammasome activation (77). Caspase 11 is activated by penta- and hexa-acylated lipid A, but not by tetra-acylated lipid A (53). As consequence, hexa-acylated lipid A triggers IL1β secretion and pyroptosis, while tetra-acylated lipid A does not (54). This strategy has been exploited by other pathogens, such as *Shigella*, which features tetra-acylated lipid A in the initial phase of infection, reducing its recognition by the immune system. In later stages of infection, the bacteria express hexa-acylated lipid A, which efficiently activates the immune system with deleterious effect (77). Similarly, LPS of *Chlamydia trachomatis* does not activate the non-canonical inflammasome pathway, which was hypothesized to depend on its unique structure (78). Chlamydial lipid A has a penta-acylated structure, due to the absence of the *lpxM* gene (79). *C. burnetii*, which lacks the enzymes LpxM and LpxL responsible for catalyzing the last steps of lipid A synthesis, produces a tetra-acylated lipid A (52). In contrast to murine caspase 11, human caspase 4 can detect tetra-acylated LPS, which contributes to different responses of human and murine cells in response to under-acylated LPS (80). Consequently, the acylation pattern of lipid A might by itself explain the lack of potent caspase 11 activation by *C. burnetii* and the difference in activation of the non-canonical inflammasome between human and murine macrophages (32–34, 67). Importantly, modulation of the acylation pattern of *C. burnetii* lipid A improved activation of the non-canonical inflammasome and reduced the bacterial burden (Fig. 10). It has to be taken into account, that the mutant mainly produced tetraacylated lipid A, while hyperacylated species (penta-and hexa-acylated lipid A) were present in minor amounts (Figs. 11 and 12). It can only be speculated why the low amount of hyperacylated lipid A is sufficient to trigger inflammasome activation and control of the infection. It might be possible that it is sufficient to reach a threshold required for activation. In addition, activation might be dose-dependent. Thus, shifting the ratio toward greater hyperacylation might further increase non-canonical inflammasome activation and other immune responses to achieve even better infection control.

In future studies it will be necessary to more precisely modulate the acylation pattern of lipid A to determine the immune response to tetra-, penta- and hexa-acylated lipid A in a murine *in vivo* model of chronic Q fever infection (81). This will allow to judge the importance of this immune evasion ability in the development of latent *C. burnetii* infection and/or the progression to chronic Q fever. However, it has to be taken into account that besides acylation of lipid A, *C. burnetii* might also use post-transcriptional modulation, such as ubiquitination, phosphorylation and acetylation, to evade host cell immune responses.

## MATERIAL and METHODS

### Reagents

Chemicals were purchased from Sigma Aldrich unless indicated otherwise.

### Mice

The C57BL/6 wild-type mice were obtained from Charles River Laboratories. The *Acod1*^+/-^ and *Acod1*^-/-^ mice (C57BL/6NJ-Acod1em1(IMPC)J/J) were generated by Jackson laboratories (Strain #:029340), obtained from Dr. Aline Bozec, and bred at the Präklinisches Experimentelles Tierzentrum of the Medical Faculty at the Friedrich Alexander Universität Erlangen-Nürnberg.

### Isolation and Differentiation of Murine Bone Marrow Derived Macrophages

Femurs and tibiae were harvested from C57BL/6 wild-type, ACOD1^+/-^, and ACOD1^-/-^ mice. The bone marrow was flushed out using a 27G syringe (BD bioscience, Germany) and centrifuged at 365xg for 10min. The resulting pellet was resuspended in macrophage differentiation medium (MDM) composed of Dulbecco’s modified Eagle’s medium + GlutaMAX (DMEM, Gibco, Germany), 15% fetal calf serum (FCS) (Sigma Aldrich, Germany), 20% supernatant from L929 cells, 1% penicillin/streptomycin (P/S, Gibco, Germany), 1% Non-essential amino acids (NEAS, Gibco, Germany) and 0.5% HEPES. Following cell counting, the cells were seeded into T75 culture flasks (Thermo Fisher Scientific, Germany) at a density of 7-10^6^ cells/flask and were differentiated for 7-9 days at 37°C and 5% CO_2_. After 4 days of differentiation, additional MDM was added to the BMDM.

### Cell culture

Differentiated BMDMs were harvested by aspirating the medium, adding cold 1x PBS (biowest, Germany), and gently scraping off the macrophages using a cell scraper. The macrophages were centrifuged at 365xg for 10min. The resulting pellet was resuspended in cMoAb (RPMI 1640 (Gibco, Germany) supplemented with 10% FCS, 1% HEPES, and 0.5% ß-mercaptoethanol) and the macrophages were plated at a density of 1x10^6^/well in a 12-well plate to adhere overnight at 37°C, 5% CO_2_.

### Culture of Coxiella burnetii

The *Coxiella burnetii* NMII and *C. burnetii* Δ*dotA* strains (18, 38) were inoculated at a concentration of 10^6^/ml into ACCM-2 (Sunrise Science Products, USA). Following a five-day incubation at 37°C, 5% CO_2_ and 2.5% O_2_, the bacterial cultures were centrifuged at 738xg for 20min. The pellets were resuspended in cMoAb and the optical density (OD) was measured at 600nm. An OD reading of one corresponds to a bacterial concentration of 10^9^/ml.

### Preparation of Legionella pneumophila ΔflaA

The *Legionella pneumophila* serogroup 1 Δ*flaA* strain (60) was cultured at 37°C on buffered charcoal yeast extract agar plates. Single colonies were picked and streaked for a two-day heavy patch (19). Bacteria were harvested and resuspended in cMoAb medium. The bacterial concentration was determined by measuring the OD_600_, with 1 OD_600_=10^9^/ml.

### Infection

Prior to infection, the medium of the macrophage culture was exchanged to RPMI supplemented with 0.1% FCS, 1% HEPES, and 0.5% ß-mercaptoethanol. BMDMs were either left uninfected or were infected at a multiplicity of infection (MOI) of 3, unless otherwise specified. BMDM were infected with *L. p. ΔflaA* for 8h or with *C. burnetii* for 32h (19). For double stimulation/infection, BMDM were first infected with *C. burnetii* for 24h, followed by additional 8h co-infection with *L. p.* Δ*flaA* or stimulation with outer membrane vesicles (OMVs) as described below. The cells were incubated either under normoxic (21% O_2_) or hypoxic (0.5% O_2_) conditions.

### Stimulation

BMDM were stimulated with 2µg/ml OMVs prepared from *E. coli* BL21, with sizes from 35 to 60nm (InvivoGen, Germany) for 8h. As a positive control, the BMDM were first primed with 1µg/ml *E. coli* LPS (Sigma Aldrich, Germany) for 4h, followed by activation with 15µM Nigericin (Sigma Aldrich, Germany) for 30–45min.

### IL1ß ELISA

The supernatants collected from uninfected, infected and OMV-treated BMDM were subjected to IL1ß secretion analysis via ELISA according to the manufactureŕs instructions (Mouse IL1 beta/IL1F2 DuoSet ELISA, R&D, Germany).

### Colony Forming Unit (CFU) Determination

The supernatant from infected BMDM was aspirated, and ice-cold ddH_2_O was added to the cells for lysis. The lysate was centrifuged at 20,817xg for 1min. The pellet was resuspended in ACCM-2 medium. For colony-forming unit (CFU) determination, a 10-fold dilution series was prepared extending up to 10^-5^ (82). 10µl of each dilution was plated on ACCM-2 agar plates (containing 0.3% agarose) in triplicates. The plates were incubated for 9-10 days at 37°C, 2.5% O_2_, and 5% CO_2_ prior to colony counting.

### RNA Extraction

Macrophages were collected, seeded at a density of 2x10^6^ cells/well in a 6-well plate, and infected/stimulated as mentioned. To extract RNA, the medium was removed, and the cells were lysed using 350µl of Lysis Buffer (39). Subsequent RNA purification was conducted following the manufacturer’s protocol, utilizing the NucleoSpin RNA Plus Mini kit for RNA purification with DNA removal columns (Macherey & Nagel, Germany).

### cDNA Synthesis and qPCR

RNA concentration was measured using a Nano-Drop (Thermo Fisher Scientific, Germany) and an RNA quantity from 500ng–1µg was used for cDNA synthesis with the High-Capacity cDNA Reverse Transcription Kit (applied biosystems, Germany). Subsequent qPCR was performed using the PowerUp SYBR Green Master Mix (applied biosystems, Germany), with a cDNA dilution of 1:10. The primers used are listed in Table 1.

**Table 1:**
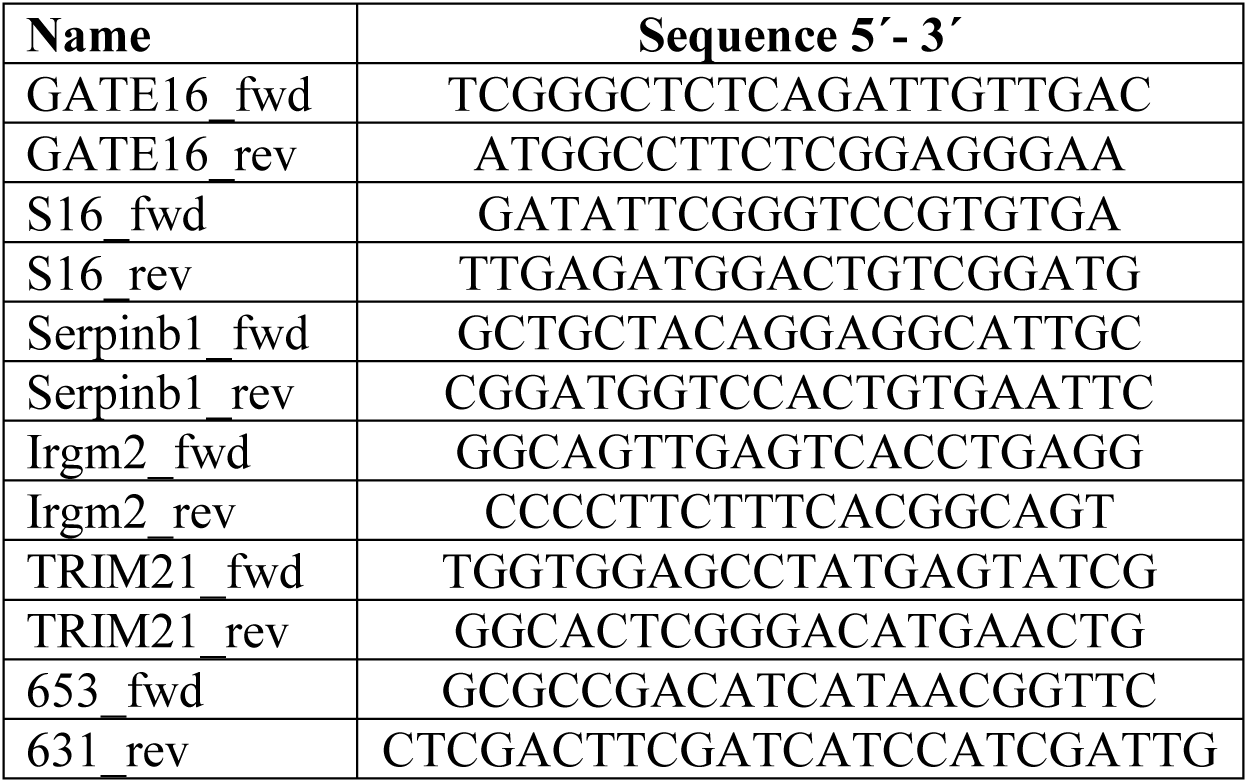
Primers used.

### Detection of caspase 11 oligomers

BMDM were either not infected, infected with *C. burnetii* for 32h, or infected first with *C. burnetii* for 24h, followed by additional 8h co-infection with *L. pneumophila* Δ*flaA* as described above. The cells were treated as previously described (29). In detail, 4x10^6^ BMDM were detached by accutase (BioLegend, USA) treatment for 15min at 37°C. Cells were pelleted by centrifugation at 365xg for 5min at 4°C and washed 3x with ice cold PBS. To stabilize formed caspase 11 oligomers, the cells were resuspended in PBS and DSS (Thermo Fisher Scientific, UK) was added to a final concentration of 500 µM. The crosslinking reaction was incubated for 30min at RT. Afterwards, the reaction was quenched by addition of 1M Tris-HCl (pH 7.5) to a final concentration of 100 mM. After stopping the reaction, the cells were washed 3x with ice cold PBS and lysed using 0.5x non-reducing LDS loading buffer (Thermo Fisher Scientific, UK). The lysates were incubated for 10min at 95°C and stored at −20°C until immunoblot analysis was performed.

### Immunoblot analysis

Macrophages were seeded at a density of 2x10^6^/well in a 6-well plate. Samples were harvested by addition of 100µl 2x Laemmli (250 mM Tris-HCl pH 6.8, 8% SDS, 40% glycerol, 8% β-mercaptoethanol, and 0.02% bromophenol blue) to the cells. The samples were sonicated for 5min and boiled for 10min. Proteins were separated by gel electrophoresis on a Bolt 4-12 % Bis-Tris Plus gel or on a 4-15 % Mini-PROTEAN TGX protein gel (Thermo Fisher Scientific, Germany) and transferred to a nitrocellulose membrane (Millipore, Germany). After blocking with 5% BSA or skimmed milk in PBS-T, the membrane was incubated overnight at 4°C with antibodies targeting GBP1 (Invitrogen, 1:1000), caspase 11, cleaved gasdermin D, NLRP3, cleaved and uncleaved IL1ß (all Cell signaling, 1:1000) or actin (Sigma Aldrich, 1:2500). As secondary antibodies, either Goat IgG anti-Rabbit IgG (H+L)-HRP (dianova, Germany, 1:5000) or Goat IgG anti-Rat IgG (L)-HRP (dianova, Germany, 1:5000) were utilized. Detection of the proteins was performed using a chemiluminescence detection system (Thermo Fisher Scientific, Germany).

### Crystal Violet Staining

BMDM were seeded at a density of 2x10^6^/well in a 6-well plate, and treated as described. The supernatant was aspirated, and the cells were rinsed with PBS. The cells were fixed with 3% paraformaldehyde for 3min. After three washing steps with PBS, a 0.2% crystal violet solution in 10% methanol was added for 10min. The cells were washed three times with PBS. Finally, a 0.5% SDS solution was added for 20min and the absorbance was measured at 550nm.

### LDH Assay

BMDM were seeded into 12-well plates at a density of 10^6^/well and were infected as described above. At the time points indicated, the culture supernatant was collected and its LDH content was measured using the CytoTox 96 Non-Radioactive Cytotoxicity Assay (Promega, Germany) according to the manufacturer’s instructions. As a positive control, the lysis buffer provided in the kit was used to determine the maximum LDH release. Absorbance was measured at 490nm.

### Cloning strategy

The constructs leading to the final *lpxM*-His-*lpxL*-HA-IGR-*lpxD*-3xFlag-*lpxA*-HA cistron were cloned into the pKM244mod vector for expression in *C. burnetii* (83). The genes *lpxM*, *lpxL*, *lpxD*, and *lpxA* derived from *E. coli* K12 MG1655 (Accession no. CP097882.1), were codon-optimized for expression in *C. burnetii*, and the complete constructs, including epitope tags (His, HA or Flag) and intergenic region (IGR), were synthesized by IDT.

Initially, *lpxM*-His-*lpxL*-HA was inserted into pKM244mod under the control of an IPTG-inducible promoter. The corresponding target DNA was amplified via PCR using Q5 Polymerase (New England Biolabs) and purified with the QIAquick PCR Purification Kit (Qiagen). Both the target DNA and pKM244mod were digested with KpnI-HF and EcoRI-HF (New England Biolabs) for 1h at 37°C, 350rpm. The digested PCR product was purified using the QIAquick PCR Purification Kit, while the vector, along with an uncut control, was separated via gel electrophoresis (1.5% agarose, 120V, 30min) and purified by gel extraction with the QIAquick PCR & Gel Cleanup Kit (Qiagen). The digested target DNA was then ligated into pKM244mod using T4 DNA Ligase (Thermo Fisher Scientific), following the manufacturer’s protocol, and incubated at RT for 1h. To introduce IGR-*lpxD*-3xFlag-*lpxA*-HA, the target DNA was similarly amplified, purified, and digested along with the plasmid using NsiI-HF and DraIII-HF (New England Biolabs). Ligation was performed as described above, resulting in a final plasmid containing the *lpxM*-His-*lpxL*-HA-*lpxD*-3xFlag-*lpxA*-HA construct (Figure S2).

### Transformation of *E. coli*

For transformation, 10µl of the ligation reaction was added to 50µl competent *E. coli* DH5α cells. The mixture was kept on ice for 30min, followed by heat shock treatment for 45s at 42°C, and chilled on ice for 2min. After adding 1ml LB medium, the culture was incubated for 1.5h at 37°C and plated on LB agar plates with ampicillin (100µg/ml) for overnight incubation at 37°C.

### Colony PCR

To confirm target gene integration, colony PCR was performed using flanking primers (653 and 631, Table 1). Single colonies were resuspended in 5µl H₂O and additionally streaked onto LB + ampicillin plates. The suspension served as the PCR template, and reactions were carried out with DreamTaq DNA Polymerase (Thermo Fisher Scientific) following the manufacturer’s protocol.

### Induction and analysis of target gene expression in *E. coli*

To assess target protein expression (60), a single *E. coli* colony was cultured overnight in LB + ampicillin (100µg/ml, 37°C). The next day, 60µl of the overnight culture was diluted 1:50 in fresh LB + ampicillin and grown to OD_600_∼0.7. The culture was then split, and either induced or not with 1mM IPTG for 3–4 h. Bacteria were pelleted (20,817xg, 1 min), resuspended in 50µl 2x Laemmli buffer and boiled for 5min). Protein expression was analyzed by immunoblot using HA- (Roche), His- (Cell Signaling)-, and Flag- (Sigma Aldrich)-tag antibodies.

### Plasmid preparation

For a DNA mini prep, a single *E. coli* colony harboring the plasmid was cultured overnight in LB medium supplemented with ampicillin (100µg/ml) at 37°C with rotation. Plasmid DNA was extracted using the NucleoSpin Plasmid Kit (Macherey-Nagel). For a maxi prep, a pre-culture in LB medium with ampicillin (100µg/ml) was grown for 8h at 37°C with shaking. Then, 100µl of the pre-culture was transferred to 2xYT medium with ampicillin for overnight cultivation (37°C, 150rpm). Plasmid DNA was extracted using the NucleoBond Xtra Midi Kit (Macherey-Nagel).

### Sequencing

To verify the target gene sequence, plasmid DNA was sequenced by Macrogen and Eurofins. Sequences were analyzed using Lasergene software.

### Electroporation and isolation of single *C. burnetii* mutant clones

*C. burnetii* cultures were incubated in ACCM-2 medium for 5 days, then centrifuged (738xg, 20min) and resuspended in 20% glycerol. After measuring the OD_600_, the bacteria were pelleted and adjusted to 5.5x10^9^/ml in 20% glycerol. For electroporation, 50µl of the bacterial suspension was mixed with > 10µg of plasmid-DNA and incubated for 10min at RT (84). The mixture was transferred to a pre-cooled 1mm cuvette and electroporated using a Bio-Rad Gene Pulser Xcell (1800V, 25µF, 500Ω). After 5min on ice, RPMI + 10% FCS was added and mixed. Subsequently, ACCM-2 was added, and the culture was incubated overnight (37°C, 2.5% O_2_, 5% CO_2_). Then, fresh ACCM-2 containing chloramphenicol (final concentration: 3µg/ml) was applied, and the culture maintained for an additional 6 days. The bacteria were plated on ACCM-2 agar plates (0.3% agarose) supplemented with 5µg/ml chloramphenicol and incubated for 10-14 days (37°C, 2.5% O_2_, 5% CO_2_). Single colonies were transferred to a 96-well plate with ACCM-2 + chloramphenicol and sequentially expanded.

### Screening for positive *C. burnetii* transformants

To confirm plasmid presence and gene expression, single colony cultures were induced with 1 mM IPTG. After 24 h, the bacteria were centrifuged (20,817 x g, 1min), the pellets resuspended in 2x Laemmli buffer, then sonicated (5min) and boiled (95°C, 10min). Protein expression was analyzed via SDS-PAGE and wet western blotting using HA-, His, or Flag-Tag antibodies.

### Cultivation and preparation of LPS-modified *C. burnetii* for LPS analysis

Wild-type and LPS-modified *C. burnetii* strains were pre-cultured in ACCM-2 for 5 days before expansion into T175 flasks. Chloramphenicol (3µg/ml) was added to modified strains for selection. Cultures were grown to OD_600_ 0.2–0.3, after which modified strains were induced with 1mM IPTG, while the wild-type remained uninduced. After 24h, the cultures were centrifuged (738xg, 20min), the pellets resuspended in PBS + 2% formaldehyde, and stored at 4°C for 14 days to ensure complete inactivation before LPS analysis.

### Cultivation of LPS-modified *C. burnetii* for infection experiments

Modified *C. burnetii* was inoculated at 10^6^/ml in ACCM-2 containing chloramphenicol (3µg/ml) and cultured for 4 days at 2.5% O_2_, 5% CO_2_ and 37°C. Expression was induced with 1mM IPTG, followed by an additional 24h-incubation. Harvesting and subsequent experimental procedures were performed as for wild-type *C. burnetii*.

### LPS isolation

*C. burnetii* cells (250mg wet pellet weight) underwent an overnight wash with chloroform-methanol (2:1, v/v) at 20°C to achieve phospholipid removal. Following centrifugation at 3000xg for 10min at 20°C, the resulting cell pellet was re-suspended in 15ml of preheated distilled water (68°C). The suspension was then extracted with an equal volume of aqueous 90% phenol (Sigma-Aldrich, Germany) according to the procedure by Westphal and Jann (85). The aqueous phase was collected after centrifugation, the phenol phase was subjected to two subsequent extractions with water. The combined aqueous phases were dialyzed against distilled water for 3 days and lyophilized.

### Acetic acid hydrolysis

LPS were hydrolyzed in 1% acetic acid (2mg/ml) at 100°C for 1.5h, followed by lyophilisation (86). Lipid A was extracted from the resulting lyophilized residue using two successive extractions according to the Matyash method with minor modifications. For the first extraction, a methyl tert-butyl ether (MTBE) /water mixture (4:1.25 v/v) was used, the second extraction involved 0.5ml of a MTBE /methanol/water mixture (10:3:2.5, v/v/v) (87). The organic phases from both extractions were collected and dried under a stream of nitrogen gas (N₂). The dried lipid A was solubilized in 50µl of chloroform.

### MALDI-TOF/TOF

MALDI-TOF analyses were performed using a Bruker UltrafleXtreme Matrix-assisted laser desorption/ionization time-of-flight mass spectrometer (MALDI-TOF/TOF, Bruker Daltonics, USA). The ion source voltage was set to 20kV in negative mode, the negative mass spectrum was scanned between m/z 700 and 3000. Norharmane (Sigma-Aldrich, Co., St. Louis, MO) was used as the matrix at a concentration of 20mg/ml, dissolved in a chloroform:methanol mixture (2:1, v/v). The samples (1µl) were mixed with 1µl of the matrix solution and spotted onto an MTP AnchorChip 384 steel target (Bruker Daltonics, USA).

### STATISTICS

Statistical analyses were conducted using GraphPad Prism (version 10), while graphical visualizations were created in R (version 4.3.2). Plots containing broken axes were produced with the R package ggbreak (88). Normality was evaluated using the Kolmogorov–Smirnov test, D’Agostino & Pearson test, or Shapiro–Wilk test. Based on the outcome of the normality assessment, either parametric or non-parametric statistical tests were applied.

## ACKNOWLEDGEMENTS

This work was supported by the Deutsche Forschungsgemeinschaft (DFG, German Research Foundation): project A06 in CRC1181 “Resolution of Inflammation” to AL and RL; project A3 within the Research Training Group “Immunomicrotope” (GRK 2740/447268119) to AL; project MIMIC to AL; project LU 1357/5-2 to AL. We thank Dr. Jan Schulze-Luehrmann with the setting up of experiments needed for the revision.

## AUTHORS CONTRIBUTION

AL conceived and designed the study with support from KMS and CB. MS, FA, FC and ES performed the experiments. MS, FC, LS, ES and AL analyzed the data. AL, RL, KMS, LS and CB provided resources. AL supervised the study. MS and AL wrote the manuscript. All authors reviewed and approved the final manuscript.

