## Supplemental for "The lipid A acylation pattern of *Coxiella burnetii* prevents detection and clearance by the non-canonical inflammasome in primary murine macrophages"

**Supplemental Information**

Figure S1

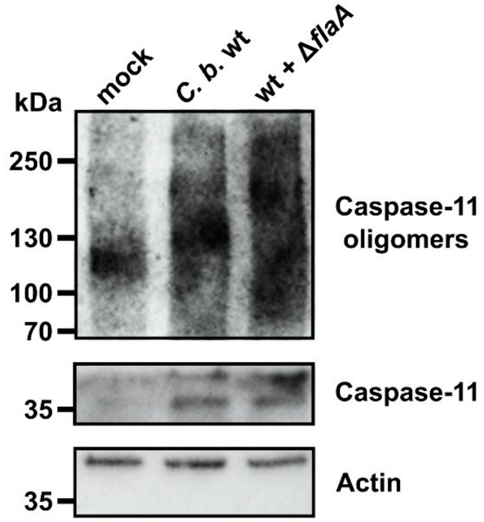


**Figure S1: Infection with *C. burnetii* fails to potently induce caspase 11 oligomerization in murine BMDM.** BMDM were either not treated or infected with *C. burnetii* wt for 32h or infected with *C. burnetii* wt for 24h with subsequent infection with *L. pneumophila* Δ*flaA* for 8h. For all infections a MOI of 3 was used. Immunoblot analysis was performed using antibodies against caspase 11 and actin as loading control. One representative blot from at least three independent experiments is shown.

Figure S2

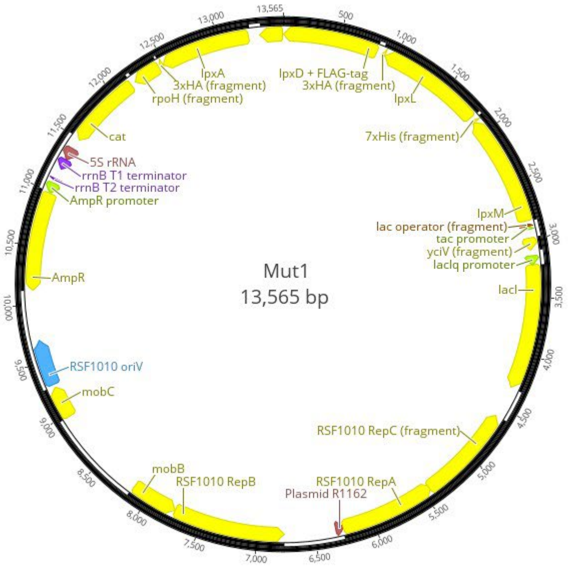


**Figure S2: Plasmid map of pKM244mod-*lpxM*-His-*lpxL*-HA-*lpxD*-3xFlag-*lpxA*-HA.** This shuttle vector contains the RSF1010 *oriV* as replication origin and ampicillin and chloramphenicol resistance cassettes for selection in *Escherichia coli* or *C. burnetii*, respectively. The expression of LpxA-HA, LpxD-Flag, LpxL-HA and LpxM-His is driven by the IPTG-inducible *tac* promotor.

**Table S1: Spectra of the MALDI-TOF mass spectrometry from figure 11.**

| m/z | S/N | Quality Fac. | Res. | Intens. | Area |
|---|---|---|---|---|---|
| 708.243 | 2.1 | 377 | 239 | 3939.28 | 19525 |
| 724.246 | 3.7 | 891 | 246 | 6909.51 | 34626 |
| 736.452 | 2.2 | 351 | 749 | 4734.35 | 9067 |
| 738.611 | 4.5 | 6505 | 487 | 9655.71 | 28175 |
| 752.776 | 2.4 | 3684 | 304 | 4408.59 | 20843 |
| 766.485 | 2.0 | 442 | 630 | 4379.50 | 10787 |
| 1126.105 | 2.6 | 4080 | 359 | 3730.49 | 26589 |
| 1140.392 | 3.2 | 2935 | 830 | 5470.34 | 18783 |
| 1144.731 | 2.4 | 2137 | 397 | 3563.92 | 22041 |
| 1155.739 | 4.2 | 614 | 1304 | 8008.76 | 16619 |
| 1158.593 | 7.5 | 3214 | 450 | 10286.36 | 63959 |

| | | | | | |
|---|---|---|---|---|---|
| 1169.423 | 3.5 | 281 | 828 | 5894.57 | 17688 |
| 1173.126 | 9.1 | 2905 | 369 | 12535.87 | 93317 |
| 1189.032 | 4.0 | 1061 | 255 | 6172.48 | 64191 |
| 1201.095 | 2.3 | 412 | 1614 | 4362.80 | 8203 |
| 1203.511 | 3.3 | 1316 | 548 | 4692.41 | 23966 |
| 1207.118 | 4.1 | 4555 | 386 | 5799.77 | 41942 |
| 1219.381 | 3.5 | 1438 | 685 | 5317.00 | 21055 |
| 1222.105 | 4.3 | 7456 | 401 | 5730.17 | 42391 |
| 1235.775 | 2.9 | 3256 | 520 | 4169.10 | 22408 |
| 1238.814 | 2.9 | 5478 | 444 | 4138.96 | 26203 |
| 1248.851 | 2.9 | 819 | 651 | 3976.88 | 18721 |
| 1252.864 | 2.6 | 1909 | 450 | 3546.47 | 22545 |
| 1309.961 | 2.3 | 3362 | 1035 | 3295.28 | 11080 |
| 1320.940 | 4.1 | 862 | 1124 | 5754.17 | 18733 |
| 1323.848 | 9.6 | 14626 | 744 | 11759.59 | 56528 |
| 1333.204 | 2.6 | 57 | 1728 | 4194.56 | 8052 |
| 1334.982 | 9.5 | 1176 | 900 | 12292.55 | 46917 |
| 1337.988 | 29.5 | 41074 | 733 | 34529.73 | 168603 |
| 1349.200 | 17.2 | 856 | 903 | 21700.22 | 82217 |
| 1352.131 | 65.1 | 62529 | 708 | 72786.03 | 370671 |
| 1363.491 | 23.8 | 496 | 1079 | 32106.42 | 102814 |
| 1366.086 | 101.7 | 52427 | 725 | 110865.38 | 577087 |
| 1377.534 | 20.8 | 494 | 1363 | 29443.30 | 78710 |
| 1380.177 | 96.1 | 71748 | 705 | 99728.38 | 536892 |
| 1389.568 | 6.9 | 99 | 2127 | 9927.89 | 16799 |
| 1391.146 | 13.1 | 1220 | 1092 | 15274.21 | 55720 |
| 1394.171 | 52.6 | 45687 | 761 | 54126.38 | 277108 |
| 1402.541 | 5.2 | 103 | 2336 | 7202.24 | 10297 |
| 1404.033 | 17.8 | 1767 | 828 | 18153.42 | 91072 |
| 1408.106 | 32.7 | 10499 | 889 | 33948.68 | 155694 |
| 1418.307 | 27.3 | 1933 | 374 | 28247.17 | 266157 |
| 1423.326 | 7.3 | 1001 | 1387 | 9623.48 | 25913 |
| 1429.329 | 5.1 | 35 | 2360 | 6856.33 | 11083 |
| 1432.030 | 86.6 | 125119 | 578 | 80779.96 | 522123 |
| 1438.460 | 2.7 | 70 | 2011 | 3575.77 | 6862 |
| 1442.698 | 8.2 | 98 | 1178 | 9543.82 | 30544 |
| 1446.123 | 126.9 | 173761 | 567 | 114287.16 | 772947 |
| 1452.982 | 4.6 | 36 | 6273 | 5952.52 | 4049 |
| 1453.869 | 6.9 | 118 | 2181 | 8963.83 | 17596 |
| 1456.648 | 9.8 | 274 | 695 | 8855.50 | 49554 |
| 1460.183 | 114.6 | 183633 | 602 | 100794.81 | 658614 |
| 1467.068 | 3.2 | 32 | 6508 | 4130.62 | 2630 |
| 1468.378 | 7.2 | 220 | 1278 | 8077.76 | 25725 |
| 1471.237 | 10.6 | 423 | 1235 | 10882.46 | 38597 |
| 1474.234 | 63.4 | 68734 | 790 | 56857.00 | 295751 |
| 1488.276 | 39.2 | 18972 | 970 | 37260.51 | 160085 |

| | | | | | |
|---|---|---|---|---|---|
| 1500.457 | 2.6 | 164 | 3585 | 3186.77 | 3912 |
| 1502.328 | 13.3 | 14020 | 1042 | 12752.08 | 51626 |
| 1534.049 | 2.0 | 362 | 556 | 1653.72 | 12152 |
| 1545.846 | 3.1 | 1629 | 1531 | 2822.52 | 8809 |
| 1548.842 | 2.5 | 2199 | 844 | 2026.27 | 10000 |
| 1561.736 | 2.5 | 3133 | 922 | 1841.61 | 10192 |
| 1600.649 | 2.1 | 611 | 603 | 1377.09 | 10399 |
| 1613.729 | 4.1 | 3011 | 712 | 2338.07 | 17037 |
| 1626.246 | 2.4 | 403 | 2598 | 2080.08 | 3954 |
| 1628.508 | 4.3 | 2746 | 875 | 2514.58 | 15551 |
| 1640.849 | 2.4 | 323 | 4874 | 2017.98 | 2149 |
| 1642.622 | 3.7 | 2951 | 1142 | 2171.11 | 10692 |
| 1657.134 | 2.6 | 1056 | 1272 | 1658.38 | 6642 |
| 1670.587 | 2.3 | 1908 | 1445 | 1382.68 | 5582 |
| 1811.301 | 3.1 | 3856 | 1217 | 1334.66 | 7534 |
| 1825.664 | 3.5 | 3524 | 1424 | 1500.17 | 7354 |
| 1840.578 | 2.0 | 843 | 1220 | 845.93 | 4357 |
